# Photoreceptor degeneration has heterogeneous effects on functional retinal ganglion cell types

**DOI:** 10.1101/2024.09.06.610955

**Authors:** Nadine Dyszkant, Jonathan Oesterle, Yongrong Qiu, Merle Harrer, Timm Schubert, Dominic Gonschorek, Thomas Euler

## Abstract

*Retinitis pigmentosa* is a hereditary disease causing progressive degeneration of rod and cone photoreceptors, with no effective therapies. Using the *rd10* mouse model, which mirrors the human condition, we examined its disease progression. Rods deteriorate by postnatal day (P) 45, followed by cone degeneration, with most photoreceptors lost by P180. Until then, retinal ganglion cells (RGCs) remain light-responsive, albeit only under photopic conditions, despite extensive outer retinal remodelling. However, it is still unknown whether the different functional RGC types remain stable, or if some types differentially alter their activity or are even lost during disease progression. Here, we addressed if and how the response diversity of functional RGC types changes with rd10 disease progression. At P30, we were able to identify all functional wild-type RGC types also in *rd10* retina, suggesting that at this early degenerative stage, the full breadth of retinal output is still present. Remarkably, we found that the fractions of functional types changed throughout progressing degeneration between *rd10* and wild-type: Responses of RGCs with ‘Off’-components (‘Off’ and ‘On-Off’ RGCs) seemed to be more vulnerable than ‘On’-cells, with ‘Fast-On’ types being the most resilient. Notably, direction-selective RGCs appeared to be more vulnerable than orientation-selective RGCs. In summary, we found differences in resilience of response types (from resilient to vulnerable): ‘Uncertain’ > ‘Fast On’ > ‘Slow On’ > ‘On-Off’ > ‘Off’. Taken together, our results suggest that *rd10* photoreceptor degeneration has heterogeneous effects on functional RGC types, with distinct sets of types losing their characteristic light responses earlier than others. This differential susceptibility of RGC circuits may be of relevance for future neuroprotective therapeutic strategies.

## Introduction

In an ageing population, an increasing number of patients suffers from blindness due to degenerative retinal diseases. For example, *Retinitis Pigmentosa* (RP), a hereditary degeneration of rod and cone photoreceptors, is highly heterogeneous with over 20 gene loci affected (reviewed in 1, 2). Consequentially, the available model systems for studying retinal degeneration (*rd*) are similarly diverse (reviewed in 3–5).

A well established model for studying the consequences of such a photoreceptor degeneration on retinal circuits is the *rd10* mouse (6). In this mutant mouse strain, a missense mutation in exon 13 of the beta sub-unit of the rod phosphodiesterase gene (Pde6*β*) triggers rod photoreceptor (rod) degeneration, which is followed by a secondary degeneration of the genetically intact cone photoreceptors (cones), eventually leading to functional blindness. Because genetic factors and likely the disease mechanisms are comparable to RP in human patients (reviewed in 7), *rd10* mice are considered a suitable model for this type of hereditary retinal dystrophy. In *rd10* mice, first signs of rod deterioration can be detected around postnatal day (P) 16, with rod loss peaking around P20, and being completed by P45 (8–10). After ∼ 6 months of age (∼ P180), the animals possess virtually no cones; thus no photoreceptors are left at this stage of degeneration in the outer retina (for overview, see Fig. 1). Typically, the degeneration starts in the mid-periphery, from where it progresses across the retina (8, 11, 12), adding a spatial axis to the degeneration.

**Fig 1.**
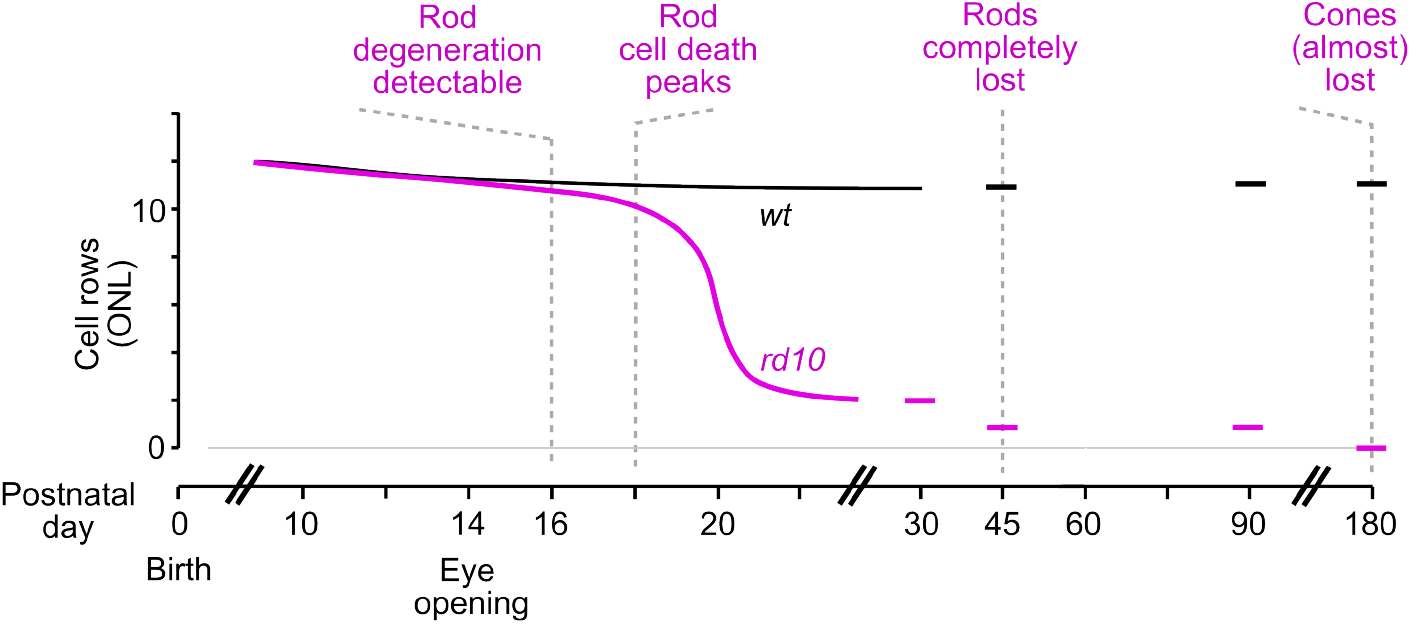
Progression of photoreceptor degeneration in the rd10 retina. Rows of cells in the outer nuclear layer (ONL) for wild-type (black) and *rd10* (magenta) retinae over time (for details and references, see text).

Due to the photoreceptor loss, the outer retina of *rd10* mice is substantially reorganised (13), resulting in new activity patterns, such as spontaneous oscillations (14). Degeneration of cells in the outer retina is followed by structural remodelling or rewiring of bipolar cells (BCs) (reviewed in 15, 16), while their postsynaptic circuits are thought to remain intact. For example, retinal ganglion cells (RGCs) seem to keep their normal dendritic morphology and projection patterns up to at least 9 weeks (∼P60), when most photoreceptors have perished (17). Accordingly, recent studies in different RP mouse models, including *rd10*, found that cones remain light-responsive and RGCs display light-evoked activity under photopic conditions until at least 2 to 3 months (∼P60) of degeneration (18, 19).

However, even if overall light-driven activity appears similar to that in healthy wild-type controls, specific RGC types still may change or even lose their functional response profile during the early phase of degeneration. This notion is supported by work showing that (partial) photoreceptor loss -experimentally initiated (20, 21) or disease-related (18, 19) affects RGC receptive fields (RFs). It is unknown, though, how the more than 40 different types of mouse RGCs (22– 24) are impacted by outer retinal degeneration. For an optic nerve crush (ONC) paradigm, it was recently shown that RGC types are differentially affected: Using transcriptomic data to investigate the survival of mouse RGC types after ONC, 25 showed that some types were more resilient and survived longer, whereas others were more vulnerable and degenerated soon after the damage. This suggests that RGC types can differ in their susceptibility to an insult.

In this study, we addressed if photoreceptor degeneration can lead to differential changes of photopic light response properties in RGCs over disease progression. To this end, we used two-photon (2P) Ca^2+^ imaging to systematically record light-evoked RGC responses in the explanted *rd10* retina and age-matched wild-type animals from P30 to P180. We found that in *rd10* a substantial number of RGCs remained light-responsive at P90 but almost none at P180, in line with earlier work (18). Notably, the RGC soma density remained virtually the same as in wild-type across the investigated age span. However, other than expected from earlier studies, we detected the first functional effects of photoreceptor degeneration on (photopic) RGC light responses already at P30, when ‘Off’-responding RGC types were less frequent in *rd10* than ‘On’-responding ones. Remarkably, other functional RGC types were affected later (e.g., ‘Fast-On’ RGCs decreased after P45), whereas the fraction of cells grouped into the ‘Uncertain’ RGCs remained largely unchanged during the investigated period.

Together, our data suggest that specific functional RGC types are more vulnerable (or resilient) than others to an insult such as photoreceptor loss. This differential susceptibility of RGC circuits may be of relevance for future neuroprotective therapeutic strategies.

## Results

To investigate how progressive photoreceptor loss affects the functional retinal output, we employed 2P Ca^2+^ population imaging in the ganglion cell layer (GCL) of *rd10* and, as a control, wild-type (C57Bl/6J) *ex vivo* mouse retina. To this end, the tissue was electroporated with the fluorescent Ca^2+^ indicator Oregon Green BAPTA-1 (OGB-1; Fig. 2A; see Methods). Here, we recorded the responses of RGCs to different light stimuli at four postnatal ages (Fig. 2B; see also Fig. 1): P30, when the retina is fully developed, rods are already significantly impacted but cones are mostly considered intact; P45, when rods are mostly lost; P90, when cones are severely damaged; and P180, when all outer-retinal photoreceptors are effectively gone (6, 8, 11, 12).

**Fig 2.**
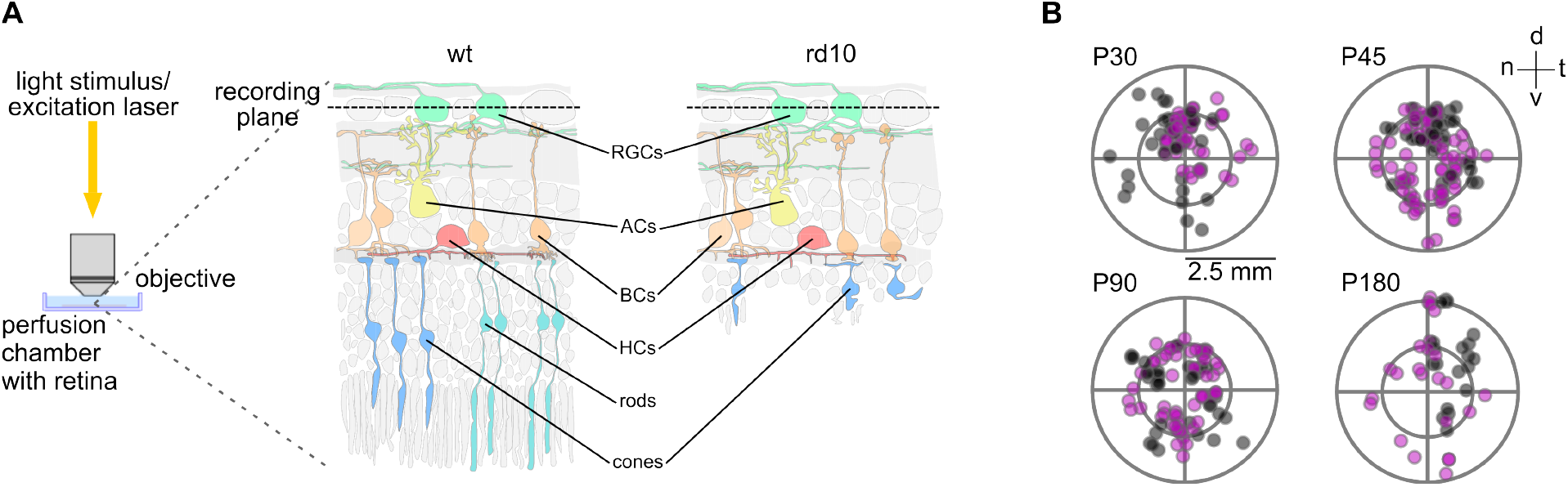
Functional Ca^2+^ imaging in *ex-vivo* wild-type and *rd10* mouse retina. (**A**) Recording configuration for two-photon (2P) imaging, with whole-mount retina in perfusion chamber and retinal ganglion cells (RGCs) facing upwards towards the objective lens (left). Illustrations of cross-sections of wild-type (wt, centre) and *rd10* (advanced degeneration, right) retina electroporated with Oregon Green BAPTA-1 (OGB-1, green). (**B**) Location of recording fields on the retina (dots) for the selected postnatal ages during degeneration in wild-type (black) and *rd10* (magenta) retinae. Retinal orientation: dorsal (d), ventral (v), nasal (n), temporal (t).

To evaluate if certain RGC types are more affected by photoreceptor degeneration than others, i.e., systematically or type-selectively, we presented a set of ‘fingerprinting’ stimuli, including full-field chirps and moving bars (Fig. 3). This allowed us to map the Ca^2+^ responses to 32 groups of RGCs (23) using a previously published RGC type classifier (26, 27, see Methods) trained on the dataset from Baden et al. (23). For simplicity, in the following, we refer to these groups as ‘functional RGC types’. While the classifier also identifies displaced amacrine cells (ACs), we focused our analyses on the RGCs.

**Fig 3.**
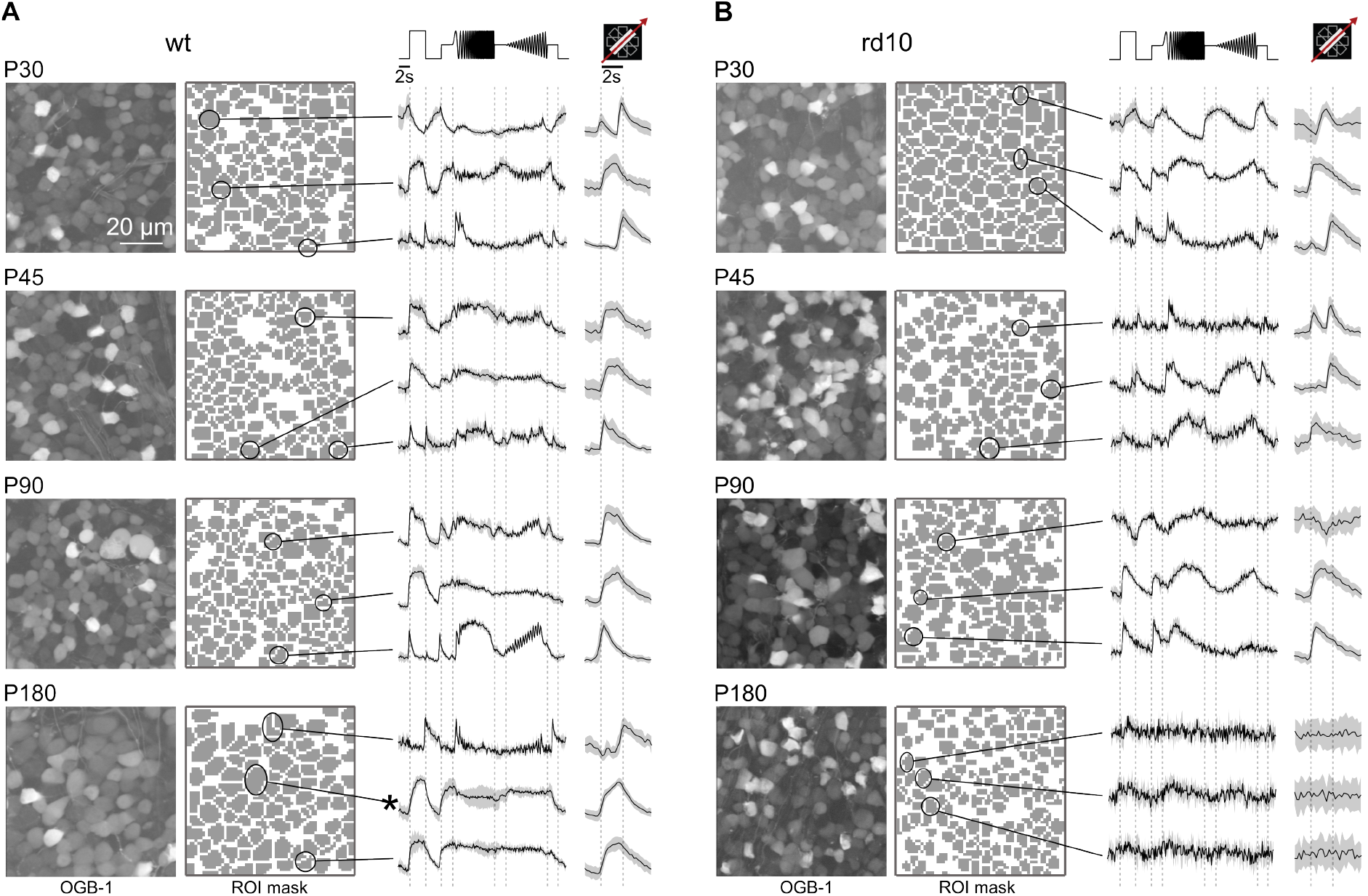
Representative recordings from wild-type and *rd10* retinae. (**A**) Recording fields in the ganglion cell layer (GCL) of wild-type retinae with cells loaded with fluorescent Ca^2+^ indicator OGB-1 (left), corresponding region-of-interest (ROI) masks (centre), and representative responses to full-field chirps and moving bar (MB) stimuli (right) of the example cells marked in the ROI masks (solid lines, mean responses; shaded areas, s.d.). Rows show examples for the four selected postnatal (P) days. Asterisk indicates alpha cell. (**B**) Like in (A) but for *rd10* retinae.

### RGC density remains constant throughout degeneration

To make sure that any difference observed between *rd10* and wild-type RGC responses is not confounded by a loss of RGCs during degeneration, we first determined the GCL soma density across postnatal ages. Previous studies have shown that RGCs, in comparison to cells in the outer retina, undergo little morphological changes during degeneration in *rd10* (e.g., 17) and, hence, we did not expected any dramatic differences in GCL soma density between the two mouse strains.

We used the semi-automatically drawn regions-of-interest (ROIs) of each 2P recording field (Fig. 3A,B; right image column) to count the number of recorded GCL cells per postnatal age and mouse line (Fig. 4A). As recording fields were of constant size (∼100 µm^2^), we used the number of cells per field as density measure (Fig. 4B). We found that GCL cell density slightly dropped after P30 in wild-type mice and then settled (P30 vs. P45, P90, P180: *p* = 0.011, 0.0008, 0.0048). This could be due to variability between fields but may also reflect the aftermath of developmental cell loss in the GCL (28). Between *rd10* and wild-type, we did not find significant differences in cell density except for P30 (*p* = 0.002). Here, the intercellular space (i.e. blood vessels) was decreased in wild-type (total white area in wild-type: *n* = 67, 429 pixels; *rd10*: *n* = 75, 682 pixels); this, in combination with general variability in the recording fields, may explain this difference at P30. In any case, our data supports that the progressive degeneration in *rd10* has no detectable effect on the overall RGC density. Additionally, we performed immunohistochemistry stainings using Calbindin antibodies, labelling most of the cells in the GCL (29). Also here, we did not observe any obvious differences in cell density, supporting our findings from functional imaging (Suppl. Fig. S1A,B, right column). Additionally, we analysed the density of alpha RGCs (30), which we could identify by their distinct large soma size (soma size *>* 136*µm*^2^, cf. example cell marked with asterisk in Fig. 3A). Again, we did not find any significant difference in density after P30 (Fig. 4C; for statistics, see legend). Support for our findings comes from additional immunolabelling experiments using SMI-32 (SMI) antibodies (see Methods; Suppl. Fig. S1A,B, left column), which strongly label alpha RGCs (31). We found that the density of SMI-positive cells in the GCL does not appear to differ between wild-type and *rd10* retinae.

**Fig 4.**
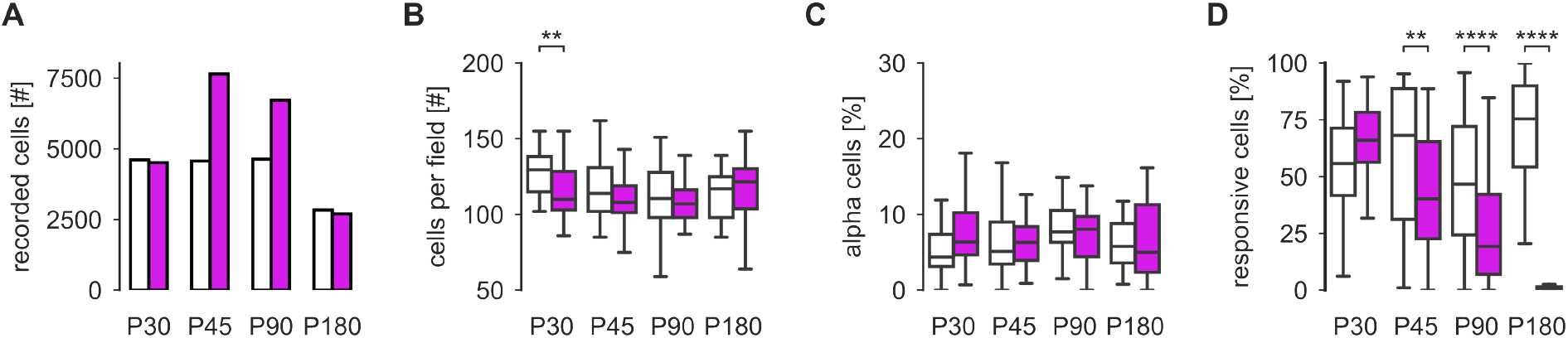
RGC density and responsiveness in wild-type and *rd10* retinae from P30 to P180. (**A**) Total number of recorded cells for each time point for wild-type (white bars) and *rd10* (magenta bars). (**B**) Average number of ganglion cell layer (GCL) somata per recording field for the different time points. (**C**) Percentage of alpha RGCs per recording field. For wild-type, the number of alpha cells differs for P90 from all other time points (vs. P30, P45, P180: *p* = 0.001, 0.019, 0.026). (**D**) Percentage of responsive cells per recording field (see Methods). For wild-type, the number of responsive cells is stable except for an increase in responsiveness between P30 and P180 (*p* = 0.027). Statistics for *rd10* RGCs: P30 vs. P45, P90, P180: *p* = 9.98 *·* 10^−5^, 2.4 *·* 10^−9^, 1.43 *·* 10^−10^; P45 vs. P90, P180: *p* = 8 *·* 10^−4^, 1.34 *·* 10^−10^; P90 vs. P180: *p* = 6.62 *·* 10^−10^. Panels (B-D): Mann-Whitney with Benjamini-Hochberg correction, asterisks indicate significance levels for comparisons between mouse lines: ∗ = *p <* 0.05, ∗∗ = *p <* 0.01, ∗ ∗ ∗∗ = *p <* 0.0001.

Taken together, our data suggest that the RGC density in *rd10* compared to wild-type does not change during the disease progression between P30 and P180.

### Responsiveness of RGCs declines with progressing degeneration

We found that the overall RGC density was unaffected by the photoreceptor degeneration (at least until P180), implying that their morphological structure may remains largely intact. To test if they also remain functional, we next measured the quality of their light-evoked responses (‘responsiveness’) from P30 to P180 (Fig. 3A,B right). To this end, we first computed response quality indices (*Qi*), which assess how reliable individual RGCs respond to the chirp and moving bar (MB) stimuli (see Methods and 23). We defined a cell as responsive if *QI*_*Chirp*_ ≥ 0.35 or *QI*_*MB*_ ≥ 0.6.

At the earliest time point (P30), the general responsiveness did not differ between wild-type and *rd10* (Fig. 4D; for all statistics, see legend). However, at P45, the responsiveness of *rd10* RGCs significantly decreased compared to wild-type RGCs. This decrease continued until P180, where the percentage of responsive *rd10* cells dropped to almost zero. In contrast, the responsiveness in wild-type did not change significantly between these ages (except for P180, 4D; for statistics, see legend), suggesting that the changes in responsiveness observed in *rd10* are mainly caused by degeneration. We also computed separately the fraction of cells that responded to each of the stimuli, using the same *QI* thresholds as before. We reasoned that the chirp and MB stimuli challenge different aspects of retinal processing: presented full-field, the chirp mainly probes temporal properties and contrast sensitivity, whereas the MB dominantly probes spatio-temporal processing (i.e. motion sensitivity, direction-selectivity).

Both *QI*_*Chirp*_ and *QI*_*MB*_ were rather constant in wild-type across ages (with a slight increase at P180), while they steadily decreased in a similar way in *rd10* (Suppl. Fig. S2A,B; for statistics, see legend). Due to the large sample sizes we also calculated the effect sizes for both wild-type and *rd10* (cf. Methods). Here, we found rather small effect sizes for wild-type, while they were medium sized in *rd10*, indicating that for wild-type significance may be mostly due to the large sample size, while in *rd10* that might not be the case. Accordingly, this was reflected in the fractions of cells responding to either stimulus (Suppl. Fig. S2C,D). At P30, the fraction of responsive *rd10* cells was somewhat higher than in wild-type, which may be related to the higher variability in the P30 wild-type data, as mentioned above.

Together, this first analysis points rather at a general, stimulus-independent decline in *rd10* RGC responsiveness.

### The majority of RGC types remains functional until late stages of degeneration

Next, we investigated if the RGC types are systematically affected by photoreceptor degeneration or if there may be type-specific differences. To this end, we employed an RGC type classifier (26) to distinguish the different functional RGC types based on their response profile to the chirp and MB stimuli, their direction-selectivity, as well as their soma size. This allowed us to map the cells onto the dataset published by Baden et al. (23) (Fig. 5). As we detected only very few responsive cells in *rd10* at P180 (Fig. 4D), we focused the further analyses on P30 to P90.

**Fig 5.**
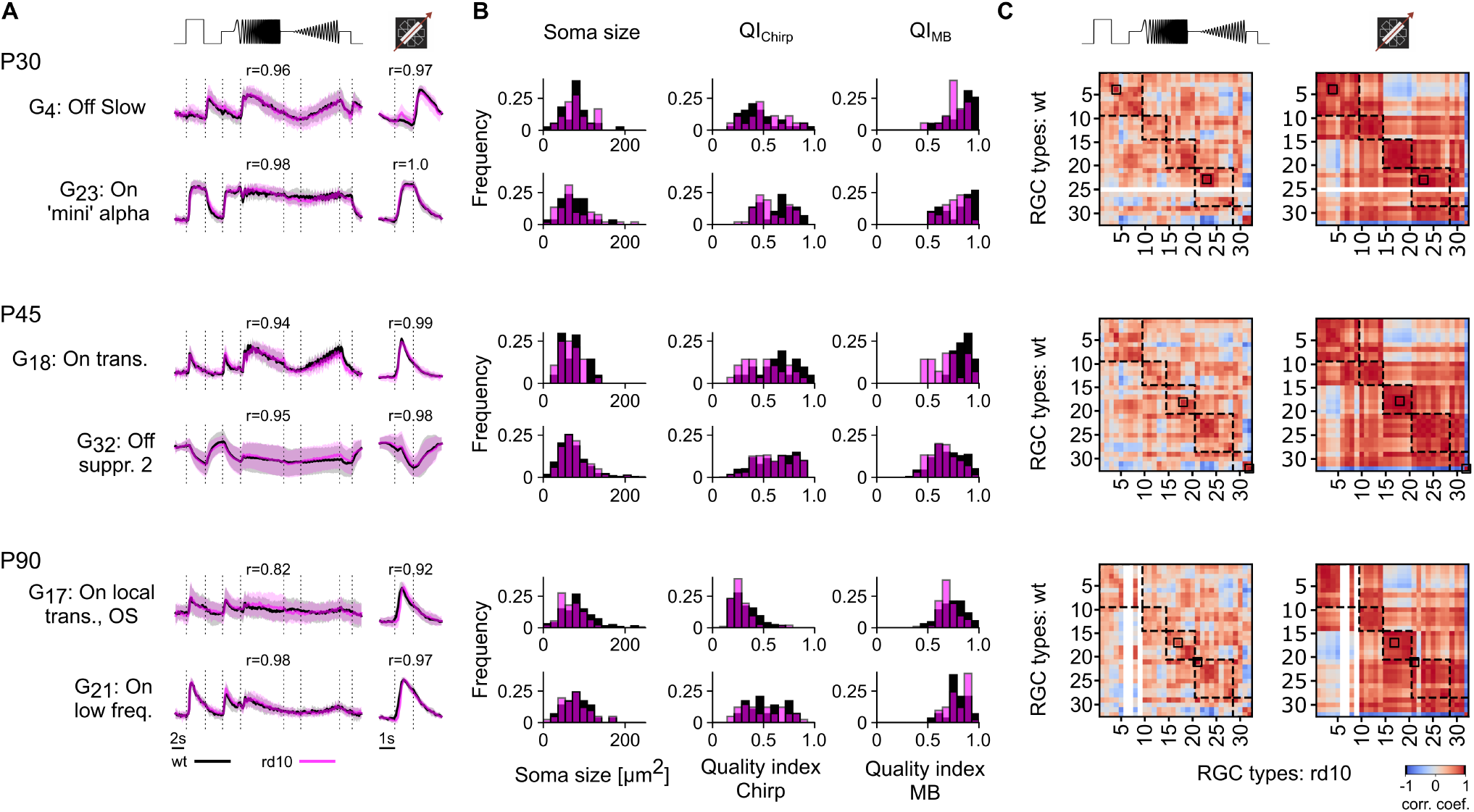
Classifying and comparing RGC responses of P30, P45 and P90, between wild-type and *rd10* retinae. **A** Representative RGC type responses to chirp (left) and moving bar (MB, right) stimuli (black, wild-type; magenta, *rd10*) for three ages (top, P30; middle, P45; bottom, P90). r-values above traces indicate Pearson correlation coefficients between the average RGC type responses in wild-type and *rd10* retinae. (**B**) Distributions of three parameters (soma size, *Qi*_*Chirp*_, *Qi*_*MB*_) of the corresponding RGC types in (A)) for *rd10* (magenta) and wild-type (black). (**C**) Correlation matrix of type average responses per RGC type between wild-type and *rd10* for chirp (left) and MB (right), with colour encoding Pearson correlation coefficient. Dashed boxes highlight the functional groups: ‘Off’, ‘On-Off’, ‘Fast On’, ‘Slow On’, and ‘Uncertain’; see (23). Black boxes within the correlation matrices indicate the corresponding RGC types from (A) White stripes indicate missing RGC types in the corresponding mouse line.

Throughout all investigated stages of degeneration, almost all of the previously identified 32 RGC types, the classifier was trained to detect, could be found not only in wildtype but also in *rd10* (31/32/32 in wild-type vs. 32/32/29 in *rd10* at P30/45/90) (Suppl. Fig. S3). Even at P90, we still found 29 types in *rd10*, despite the stark decline in responsive cells (Fig. 4D). Notably, the classifier was able to identify similar percentages of cell types across ages and mouse lines, suggesting that cell type assignment was equally reliable for both datasets (Fig. S4).

To investigate if the photoreceptor degeneration affects the response profiles of distinct RGC types, we computed the correlations of their average responses to the chirp and MB between wild-type and *rd10* at each age (Fig. 5A,C) We used this correlation as a proxy for the classifier’s ability to identify the known RGC types in *rd10* and, hence, as a measure for how well their responses were preserved. We found that the correlations between RGC response types in wild-type and *rd10* did not differ substantially across ages (Fig. 5C; Suppl. Fig. S5). However, distinct RGC types begin to disappear at P90 (cf. white vertical columns in Fig. 5C). For each classified RGC type, we also compared their soma size, *QI*_*Chirp*_, and *QI*_*MB*_ between wild-type and *rd10* and found that, overall, they matched well (Fig. 5B; see also Suppl. Fig. S6), which implies that the progressing degeneration may not introduce strong functional differences between the RGC types of both mouse strains.

Finally, we compared the correlation for each RGC type within the two mouse lines. Here, we found that it generally varies somewhat over the three ages, suggesting that there may be intrinsic variability in both mouse lines, which is potentially not related to degeneration (Suppl. Fig. S5A).

Note that a classifier-based approach cannot discover potentially new degeneration-related response types (see Discussion). Still, our type-resolved analysis clearly suggests that the functional RGC diversity in *rd10* is largely comparable to that in wild-type at least until P90. Our findings highlight the robustness of RGC diversity in *rd10* despite degeneration.

### Receptive fields tend to be smaller and have faster kinetics in *rd10*

Photoreceptor degeneration has been reported to affect RGC receptive field (RF) properties, such as RF size (18, 19, 32). Therefore, we estimated spatiotemporal RFs in both mouse lines using the responses to a shifted dense noise stimulus (33), which allowed us to probe the cells’ spatial and temporal properties in more detail. To this end, we used generalised linear models (GLMs; see Methods). As a quality measure, we only used RFs that passed the three quality thresholds: *QI*_*SVD*_ *>* 0.5, *QI*_*sRF*_ *>* 0.5 and *QI*_*tRF*_ *>* 0.85 (for details, see Methods).

We found good-quality RFs for both wild-type and *rd10* RGCs at all three time points, with the number of RFs declining in *rd10* at P90 (Suppl. Fig. S7). This decline may be at least partially cell type-specific, as for some functional types (e.g., ‘On-Off local’, G_11_), we did not find RFs at P90 in *rd10* (Fig. 6A).

**Fig 6.**
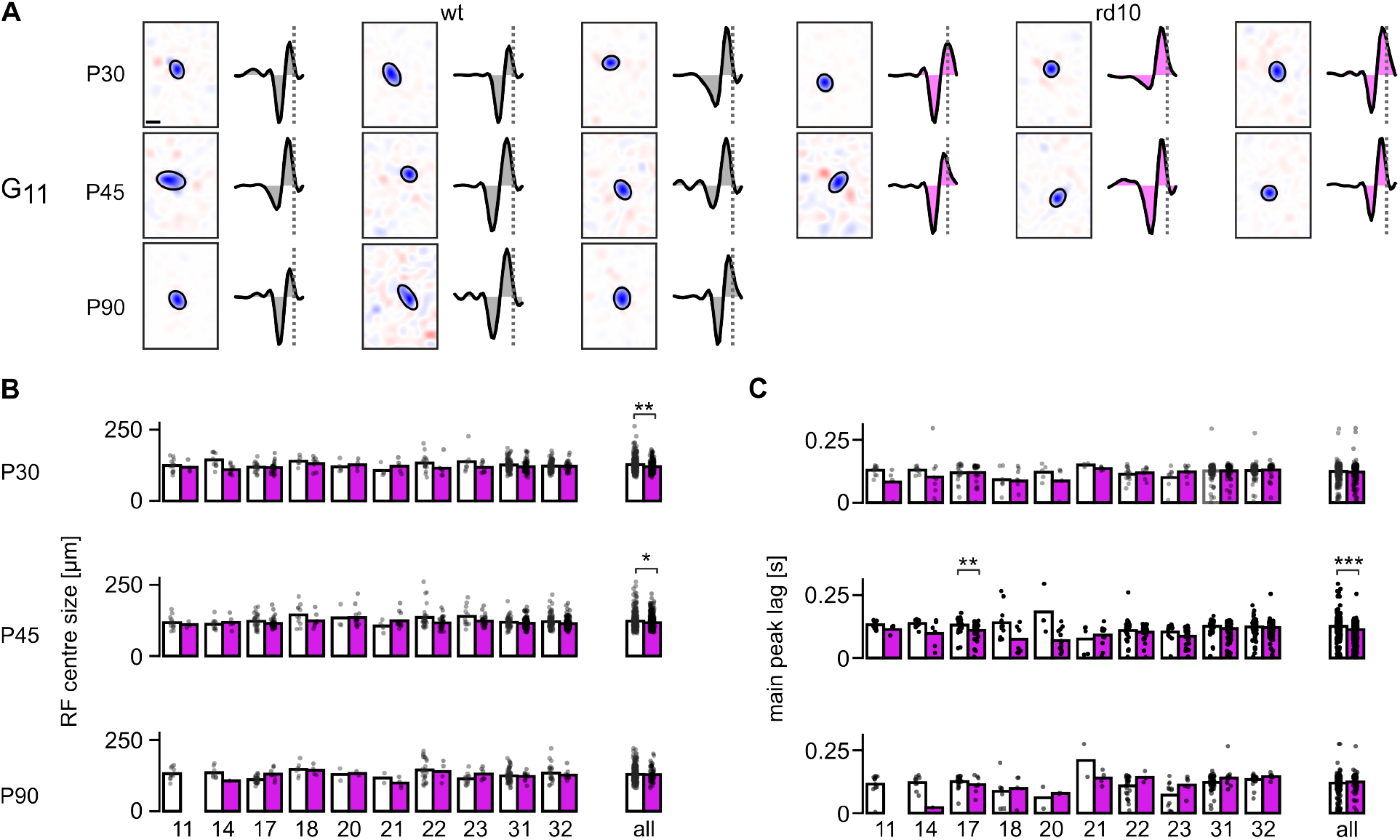
Receptive field properties in *rd10* vs. wild-type retina. (**A**) Representative spatial and temporal RFs (sRF and tRFs, respectively) of ‘On-Off local’ (G_11_) RGCs for different ages (rows) and mouse lines (wild-type, grey, left; *rd10*, magenta, right). sRF centre indicated by Gaussian fit (ellipse); scale bar: 50*µ*m; dotted line indicates onset of Ca^2+^ event in tRF, scale bar: 0.5 s. (**B**) Left: Centre size of sRF for different RGC types in wild-type (grey) and *rd10* (magenta) as mean (bar) and for individual cells (dots). Only types with *N*≥ 4 responsive cells per mouse line at P30 are shown. Right: Centre size of of all RGC types (including types not shown on the left) as mean (bar) and for all cells (dots). (**C**) Same as (B) but for main peak lag of the tRF (i.e. time to Ca^2+^ event). Panels (B-C): Mann-Whitney. On the RGC type level Benjamini-Hochberg correction was used. Tests were only calculated for *N* ≥ 4 in both groups, asterisks indicate significance levels: ∗ *<* 0.05, ∗ *<* 0.01, and ∗ ∗ ∗ *<* 0.005.

First, we investigated the size of spatial RFs (sRFs) and found that *rd10* RF centres tend to be smaller at the cell type-level, adding up to statistically smaller RFs on the population level at P30 (*rd10*: 120.18 *µ*m, wild-type: 126.92 *µ*m, *p* = 0.0014) and P45 (*rd10*: 117.2 *µ*m, wild-type: 122.5 *µ*m, *p* = 0.013), however not at P90 (*rd10*: 128.9 *µ*m, wild-type: 129.2 *µ*m, *p* = 0.829; Fig. 6B).

Next, we quantified the temporal RF (tRF) properties by computing the time lag to the main Ca^2+^ event (Δ*t* of the kernel peak closest to zero; see Methods), as a measure of tRF kinetics. This analysis revealed that overall, tRFs tended to be faster in *rd10* compared to wild-type. At P45, this difference was significant at the population level (*rd10*: 0.11 s; wild-type: 0.13 s, *p* = 4 · 10^−8^; Fig. 6C right, centre). At the cell-type level, these differences were not statistically significant except for ‘On local transient’ (G_17_) RGCs, which showed slower kinetics in *rd10* vs. wild-type at P45 (0.13 s vs. 0.11 s, *p* = 0.082) but not at P30 (0.12 s vs 0.12 s, *p* = 0.779) and P90 (0.113 s vs 0.126 s, *p* = 0.5043; Fig. 6C left).

Taken together, our RF analyses showed cell typespecific differences in sRFs and tRFs between *rd10* and wildtype mice. While these differences were quite heterogeneous and time points-dependent, we found overall *rd10* RFs to be smaller, especially earlier in degeneration (P30 and P45), and at least at P45, featuring faster kinetics.

### Progressive, type-specific degeneration of RGCs in the *rd10* retina

So far our analyses showed that the majority of RGC types can be found in both mouse lines at least until P90, suggesting that the overall functional diversity remains largely unaffected in *rd10*. On the other hand, we found changes in RF properties early in the degeneration. Therefore, we next asked if the relative composition of RGC types is different in *rd10* compared to wild-type (Fig. 7).

**Fig 7.**
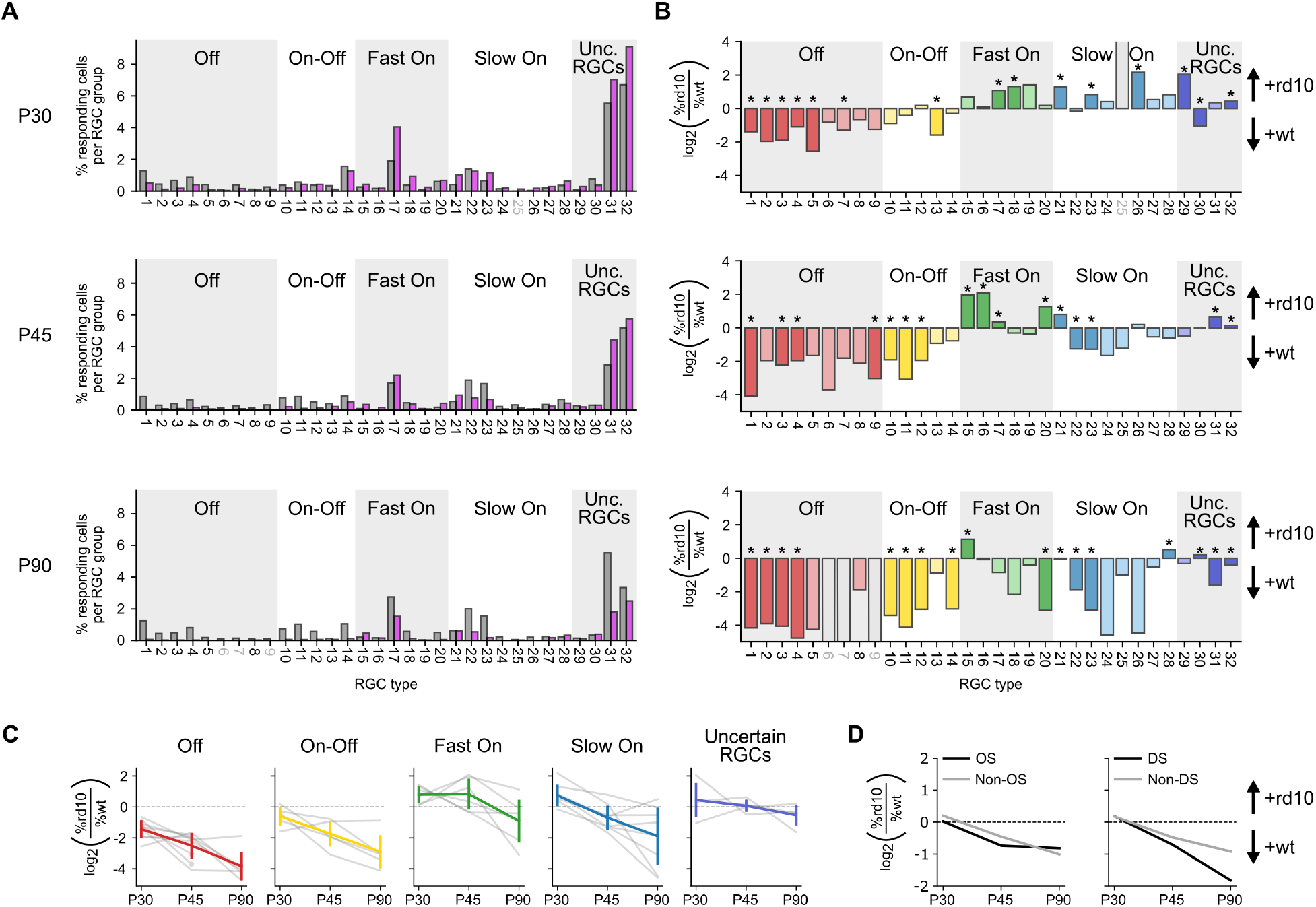
Progressive, type-specific degeneration in the *rd10* retina. (**A**) Distribution of responsive cells per functional RGC type in percent for wild-type (grey) and *rd10* (magenta) across three different ages (from top to bottom: P30, P45, and P90). (**B**) Relative abundance of functional RGC types in *rd10* vs. wild-type, quantified as *log*_2_ (*rd*10*/wt*) for the three ages. Colours indicate functional groups (‘Off’, ‘On-Off’, ‘Fast On’, ‘Slow On’, ‘Uncertain RGCs’). Positive values indicate higher abundance in *rd10*, negative values a higher abundance in wild-type. Significant differences in cell type ratios between *rd10* and wild-type (*p <* 0.01; binomial test) are marked with asterisks; Non-significant RGC types are coloured in muted colours. Open bars indicate missing types in either mouse line. (**C**) Relative abundance for the five functional RGC groups (as in (B)) as a function of age (grey, individual RGC types; coloured, mean *±* s.d.). (**D**) Relative abundance for orientation-selective (OS) and Non-OS (left) and direction-selective (DS) and Non-DS RGC types (right) as a function of age.

**Table 1.**
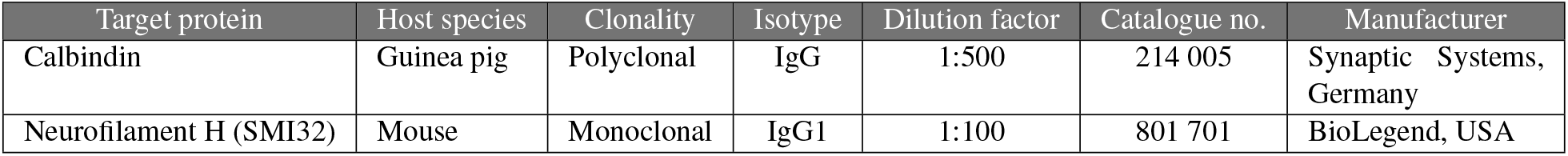
The following primary antibodies were used.

**Table 2.**
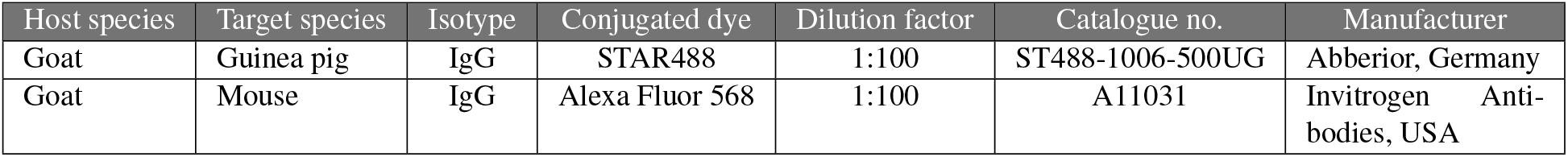
The following secondary antibodies were used.

We compared the percentage of responsive cells of all recorded cells per RGC type across mouse lines and ages (Fig. 7A). This confirmed that nearly all RGC types were present in wild-type and *rd10*, and match the response patterns reported by Baden et al. (23) quite well (Suppl. Fig. S5B,C). However, we were surprised to see that some RGC types were underrepresented (e.g., G_1,4_), while others were overrepresented (e.g., G_17,32_) in *rd10* vs. wildtype. To better visualise these differences, we calculated an index that captures the “relative abundance” of RGC types (*log*_2_(*rd*10*/wt*); Fig. 7B). This measure has previously been introduced to quantify genetic RGC types regarding their resilience to injury (25) and the contribution of RGC types to dLGN responses (34). In our study, positive values indicate higher, negative values lower abundance in *rd10*, suggestive of more resilient or more vulnerable functional types, respectively.

To our surprise, the distribution in *rd10* deviated significantly from that in wild-type as early as P30 (Fig. 7B, top). Compared to wild-type, we found fewer ‘Off’ and ‘OnOff’ cells in *rd10*, whereas the relative fraction of ‘On’ RGCs (both ‘Fast On’ and ‘Slow On’) and distinct ‘Uncertain’ RGCs (i.e. G_29_) was larger in *rd10*. At P45, most of the ‘Slow On’ cell types were less abundant in *rd10* (Fig. 7B, centre), with the exception of G_21_ (‘On low frequency’), which was more frequent. Like at P30, the ‘Fast On’ RGCs tended to be more abundant in *rd10*; this changed at P90, when only G_15_ (‘On step’) was more frequent among the ‘Fast On’ RGCs in *rd10* (Fig. 7B, bottom).

Notably, the relative abundance of RGCs with a similar response type (i.e. functional groups) changed similarly over the course of degeneration (Fig. 7C). The relative abundance became negative for ‘Off’ and somewhat less so for ‘On-Off’ RGCs already at P30, whereas for ‘Slow On’ cells, it dropped below zero for P45 and later. For ‘Fast On’ RGCs, the relative abundance became negative only at P90, while that for ‘Uncertain’ RGCs stayed close to zero for all ages. This suggests differences in resilience among functional RGC groups (from resilient to vulnerable): ‘Uncertain’ *>* ‘Fast On’ *>* ‘Slow On’ *>* ‘On-Off’ *>* ‘On’.

Finally, we evaluated if the photoreceptor degeneration affected orientation- and direction-selectivity (OS, DS; Fig. 7D). To this end, we divided RGC types into OS/Non-OS (Fig. 7D, left) and DS/Non-DS (Fig. 7D, right; see Methods). We found that the relative abundance of OS and Non-OS cells declined similarly, indicating that the loss in OS responses may simply reflect the general loss in responsiveness of OS RGCs. In contrast, the relative abundance of DS cells dropped faster than that of Non-DS cells, which may be related to the ‘On-Off’ RGCs being more susceptible to the degeneration (cf. Fig. 7B,C).

Taken together these results indicate that functional RGC types may undergo a progressive, type-specific degeneration in *rd10*.

## Discussion

Learning how degenerative diseases differentially affect functional cell types will provide a greater understanding of disease progression and may point toward novel therapeutic approaches. In this study, we measured light-evoked responses RGCs in the *rd10* mouse from P30 – when the retina is adult, rods are already impacted but cones are intact – to virtually complete photoreceptor loss at P180. We found that during this time window, the overall number of cells in the GCL was not affected by photoreceptor degeneration and comparable with the wild-type situation. To investigate how retinal output changes during disease progression, we used 2P Ca^2+^ imaging to record RGC responses to various visual stimuli. Here, we found that almost all known functional RGC types can be identified even at a late degeneration state (P90), though with slight changes in RF properties in *rd10*. Critically, however, the relative abundance of specific RGC types decreased in *rd10* vs. wild-type in a degeneration stage-dependent manner, starting with a significant loss of ‘Off’ RGC types already at P30.

### *rd10* as a model for photoreceptor degeneration in RP

Photoreceptor degeneration in the *rd10* mouse is caused by a missense mutation in the *β*-subunit of the phosphodiesterase gene (35). Rod degeneration begins around P16, shortly after eye opening (P14), with the peak occurring around P20. By P45, nearly all rods are lost, and secondary cone degeneration has begun. At least until P60, cones can still generate light-evoked responses and drive their circuitry at photopic light levels (18). Around P90, the remaining cones are severely deformed, and by P180, they are almost completely lost (for an overview, see Fig. 1, and see 8, 9, 11).

To some extend, *rd* mouse mutants, such as *rd10*, mimic human forms of autosomal recessive RP (6, 35), making them suitable for studying this condition. For example, many *rd* mice exhibit spontaneous oscillatory activity after rod loss (14, 36–38), altering the signal-to-noise ratio (SNR), and complicating therapeutic interventions (e.g., 39). Notably, these oscillations may be related to photopsia, a phenomenon observed in human RP patients, which hampers their quality of life (40; reviewed in 41).

One advantage of *rd10* vs. models like *rd1* (42) is the slower disease progression, which is more similar to that in human but still allows experimental studies in a reasonable time frame. On the other hand, RP typically starts between teenage years and young adulthood (43, 44), while in *rd10*, retinal development is not concluded when photoreceptor degeneration starts. This is most notable from the fact that *rd10* electroretinograms (ERGs) are never normal (6), already starting with decreased sensitivity at low light levels (12). Newer mouse models, such as *Cngb*1^*neo/neo*^ (45), exhibit a slower progression, which may mimic more closely the time course of human RP.

Another difference between *rd* mice and humans is the spatial gradient of degeneration: while human retinal degeneration typically starts peripherally and progresses toward the centre, leading to the initial loss of peripheral vision (43), degeneration in *rd10* mice begins more centrally (11, 12).

In summary, the *rd10* mouse model offers significant advantages for studying retinal degeneration, particularly due to its genetic relevance and well-documented disease progression. However, its limitations, such as differences in the spatial disease progression (i.e. in the context that mice lack a fovea) and the overlap between retinal development and degeneration onset, must be considered when interpreting results.

### Effects of photoreceptor degeneration on retinal organisation

Photoreceptor degeneration goes hand in hand with extensive remodelling in the outer retina of *rd* mice (13). This remodelling occurs particularly at the level of bipolar cells (BCs). Rod bipolar cells (RBCs) form new contacts with remnant cones after losing their synaptic input (46), while cone bipolar cells (CBCs) extend their branches to establish targeted connections with surviving cones after losing their original synaptic partners (47). However, these structural changes do not seem to include RGCs, with their synaptic organisation, central projections, and numbers being comparable to wild-type (17). For instance, even as their synaptic partners, type 6 CBCs, alter their dendritic connectivity, ‘On-alpha’ RGCs do not change their wiring or function (20). Still, it stands to reason that the inner retina eventually changes with progressing degeneration.

Indeed, recent studies on photoreceptor loss in *rd* mice have found subtle changes in RGCs. For example, RFs shrink, and firing rates as well as information rates decrease, in particular at low light levels (18, 19). In line with these results, we found *rd10* RFs to be smaller, partially with faster kinetics compared to wild-type. Similar reductions in RF sizes were reported in a rat RP model, though, other than our findings, RF kinetics became slower in a somewhat typespecific manner (48). Generally, changes in RFs seem to be strongly model- and insult-dependent. For example, regarding different types of alpha RGCs, partial ablation of cones, on the one hand, resulted in ‘On-alpha sustained’ RGCs displaying larger RF sizes and slower kinetics (20). A study of induced ocular hypertension, on the other hand, found ‘Off-alpha transient’ but not ‘On-alpha sustained’ cells showing a decrease in RF size (49).

These studies, together with our results, suggest that RGCs retain their functionality late into degeneration. At the same time, RGCs are affected by the remodelling – as we observed, for example, nuanced effects on temporal and spatial processing. Importantly, we found these differences to be cell type-specific and time point-dependent, as detailed in the next section.

### Differential degeneration of functional RGC types

How disease and degeneration (i.e. photoreceptor loss, glaucoma, and optic nerve injury) affect RGC health and function, in particular of specific types, has been the focus of an increasing number of studies in the past years (e.g., 20, 21, 25, 49). Since they are morphologically easy to identify, alpha RGCs have been thoroughly studied. Generally, alpha cells are considered resilient to an insult such as optic nerve crush (ONC) (50). However, a recent ONC study found differences among alpha cells, with sustained alpha cells being more resilient than transient alpha cells (25). Similarly, in a transient ocular hypertension paradigm, a glaucoma model, ‘Off-alpha transient’ RGCs were the most vulnerable of the alpha types (49). In our study, we found all ‘Off-alpha’ RGC (G_5,8_) responses to be less abundant in *rd10* vs. wild-type, whereas ‘On-alpha’ RGC (G_19,24_) responses persisted longer. This suggests that for photoreceptor loss as the insult, the divide is not between sustained and transient but rather between ‘On-’ and ‘Off-alpha’ cells, with ‘On-alpha’ cells being more resilient than their Off counterparts.

Generally, our data promotes the idea that RGC resilience in photoreceptor degeneration depends on the retinal circuits involved (e.g., ‘On’-vs. ‘Off’-pathway): Already shortly after the onset of degeneration (P30), we found the fraction of ‘Off’- and ‘On-Off’-type RGCs to be reduced, whereas ‘On’-RGCs remained functional until at least P45. Based on our data, the resilience of RGC response types can be ranked (from resilient to vulnerable): ‘Uncertain’ > ‘Fast On’ > ‘Slow On’ > ‘On-Off’ > ‘Off’. This ranking is different from that for ONC, which was studied at the RGC type-level with transcriptomic data (25), and where, for instance, no differences between ‘On’- and ‘Off’-RGCs were detected.

The difference between ‘On’ and ‘Off’ we observed is likely not directly due to changes in BC input, since in *rd10* ‘Off’-CBCs retain their function, while ‘On’-CBC function is altered (9). Instead, a recent *rd10* study using voltage-clamp recordings found an imbalance of inhibition and excitation between the ‘On’- and the ‘Off’-pathways (51). Specifically, ‘On’-type RGCs received mostly excitation, while ‘Off’-type RGCs received mostly inhibition. Carleton and Oesch (51) also established that the inhibition in the ‘Off’-pathway derives from glycinergic ACs and is stronger in *rd10* than in wild-type retina. Interestingly, this is in line with previous work on ocular hypertension, where the ‘Off’-sublamina of the IPL was demonstrated to less ‘stable’, displaying signs of disorganisation earlier than the ‘On’-sublamina (49). Moreover, the ‘Off’-stratifying dendrites of ‘On-Off’-RGCs lost their branching complexity more rapidly than the ‘On’-stratifying dendrites, while the excitatory synapses onto both ‘On’ and ‘Off’ dendrites decrease similarly. Whether a comparable change in inner retinal synaptic organisation also happens in *rd10* remains to be tested in future studies.

The differential loss of ‘Off’-type responses we observed in *rd10* is consistent with increased inhibition of the ‘Off’-pathway, possibly involving synaptic reorganisation in the inner retina (as detailed above). However, other disease-triggered mechanisms likely also play a role. These mechanisms include changes gene expression, neurotrophic factors, and Müller glia activation. For example, that RBCs switch their synaptic inputs to cones may trigger new gene expression patterns of glutamate receptors (52). Furthermore, BCs rely on neurotrophic signals from photoreceptors to maintain their dendritic arbours; the absence of these factors in the *rd* condition leads to atrophy (53, 54). Differential gradients of neurotrophic factors might also help preserve the functional layering of the IPL (49). Finally, Müller cell activation is significantly up-regulated shortly after rod loss is peaking (11). It is therefore conceivable that these glia cells become heavily involved in clearing rod remnants, which could reduce their capacity to regulate neurotransmitter uptake and maintain ion homeostasis. In turn, heightened metabolic demands and altered glial function could contribute to the distinct vulnerabilities observed in ‘Off’-pathways, which were shown to be metabolically more demanding (55).

Taken together, we propose that the resilience of RGCs is surprisingly heterogeneous, depending not only on the genetic or functional response type, but also on the type of insult to the tissue.

### Identifying cell types over the course of a progressive disease

To identify known functional RGC types, we utilised a published classifier (26) that matched our data to an established dataset (23). Specifically, this approach allowed us to estimate the probability (confidence score, see Methods) with which a cell in our wild-type and *rd10* datasets belongs to a distinct RGC type and, hence, enabled a type-resolved analysis. This analysis suggests, first, that the functional RGC diversity in *rd10* is robust and comparable to that in wild-type at least until P90, and second, a differential, RGC-type-dependent loss in responsiveness. Still, this method has its limitations: As the classifier was trained with responses from healthy wild-type retina, it cannot identify “novel” response patterns (i.e. out-of-distribution data) that may develop during disease progression. As a consequence, such novel response types will be excluded from the analysis due to their low confidence score. However, because the percentages of classified RGC types were similar across ages and mouse lines, we consider it unlikely that we missed many novel response types in *rd10*. Nevertheless, in future studies it will be important to test the possibility of an RGC type significantly changing its response properties, for example, by combining functional and transcriptomic methods in the same tissue.

### *rd10* and vision restoration

To develop successful vision restoration approaches, a better knowledge of the inner retinal circuits during disease progression is crucial. Notably, different types of pathologies (e.g., RP, glaucoma) appear to lead to differential degeneration of the RGCs. Especially in a genetically highly heterogeneous disease like RP, where a general restoration strategy may build upon the relative stability of the inner retina, it is important to know which pathways are more resilient than others. Our data suggests that interventions, such as the expression of light-sensitive channels in BCs, may be more efficient in the ‘On’-pathways, which turned out to be more resilient in *rd10*. Regarding the time point, in *rd10* mice P45 may be most suitable, when rods are mostly lost but functional cones still persist. This time-point is roughly equivalent to that when most patients are diagnosed with RP: when their night vision is heavily impaired if not lost and daylight vision is still functional (43, 44). At later time points, therapeutic approaches are likely to be severely hampered by increasing (functional) remodelling of inner retinal circuits.

## ACKNOWLEDGEMENTS

We thank François Paquet-Durand for lively and inspiring discussions, and Olga Oleksiuk for excellent microscopy support. We acknowledge support by the Open Access Publishing Fund of the University of Tübingen. This work was funded by the Tistou & Charlotte Kerstan Stiftung (RI-FG P3 EU/SCH 1-2&3 Dysz) and the German Research Foundation (DFG; SFB 1233 “Robust Vision”, 276693517; EU 42/10-1; EU 42/12-1).

## Methods

### Animals and tissue preparation

All animal experiments were conducted at the University of Tübingen. They were approved by the institutional animal welfare committee of the University of Tübingen and performed according to the laws governing animal experimentation issued by the German Government. We used retinae from C57Bl/6J (wild-type, JAX 000664) and Pde6brd10 (*rd10*, JAX 004297) mice of either sex at postnatal days (P) 30, 45, 90 and 180 (± 3 days). For recording experiments, we used *N* = 23 wild-type animals (*N* = 36 eyes) and *N* = 25 *rd10* animals (*N* = 37 eyes). For our immunohistochemistry stainings we used *N* = 10 wild-type animals (*N* = 20 eyes) and *N* = 13 *rd10* animals (*N* = 26 eyes). All animals were housed in the local animal facility under the standard 12h/12h day/night cycle at 22°C and a humidity of 55%.

The following procedures were carried out under very dim red (*>* 650 nm) light. Before each experiment, the animal was dark-adapted for *>* 1 h, then anaesthetised with isoflurane (CP-Pharma), and sacrificed by cervical dislocation. Immediately after, the eyes were enucleated with a dorsal cut as orientation landmark and hemisected in carboxygenated (95% O_2_, 5% CO_2_) artificial cerebrospinal fluid (ACSF) solution containing (in mM): 125 NaCl, 2.5 KCl, 2 CaCl_2_, 1 MgCl_2_, 1.25 NaH_2_PO_4_, 26 NaHCO_3_, 20 glucose, and 0.5 L-glutamine at pH 7.4. Sulforhodamine-101 (SR101, ≈ 0.1 *µ*M; Invitrogen) was added to the ACSF to visualise blood vessels for orientation and damaged cells in the red fluorescence channel of the microscope (56). The recording chamber was constantly perfused with carboxygenated ACSF at 4 ml/min and kept at ≈ 36°C during the experiment.

After dissection, retinae were flat-mounted on thin ceramic discs with the GCL facing up (Anodisc^*T M*^ #13, 0.1 *µ*m pore size, 13 mm in diameter, Cytiv) and bulk-electroporated with the synthetic fluorescent Ca^2+^ indicator Oregon-Green 488 BAPTA-1 (OGB-1; hexapotassium salt; Life Technologies).

To electroporate the retina (57), the anodisc was then placed between two 4 mm horizontal platinum disk electrodes (CUY700P4E/L, Nepagene/Xceltis). The lower electrode was covered with 15 *µ*l of ACSF, while a 10 *µ*l drop of 5 mM OGB-1 dissolved in ACSF covered the upper electrode and was lowered onto the tissue. Then, 9 electrical pulses (≈9.2 V, 100 ms pulse width, at 1 Hz) from a pulse generator/wide-band amplifier combination (TGP110 and WA301, Thurlby handar/Farnell) were applied to introduce the Ca^2+^ indicator into the retinal cells. Afterwards, the retina was placed in the recording chamber, with the dorsal edge of the retina pointing away from the experimenter.

The retina was left there for ≈ 30 min to recover, as well as adapted to the light stimulation by displaying a binary dense noise stimulus (20 × 15 matrix, 40 *µ*m^2^ pixels, balanced random sequence) at 5 Hz before the recordings started.

### Two-photon Ca^2+^ imaging

We used a MOM-type twophoton (2P) microscope (designed by W. Denk, MPI Heidelberg; purchased from Sutter Instruments/Science Products; 56, 58). Briefly, the system was equipped with a mode-locked Ti:Sapphire laser (Maitai-HP DeepSee, Newport Spectra-Physics) tuned to 927 nm, two fluorescence detection channels for OGB-1 (HQ 510/84, AHF/Chroma) and SR101 (HQ 630/60, AHF), and a water immersion objective (CF175 LWD×16/0.8W, DIC N2, Nikon, Germany). For image acquisition, custom-made software (ScanM by M. Müller and T.E.) running under Igor Pro 6.3 for Windows (Wavemetrics) was used, taking time-lapsed 64× 64 pixel image scans (≈ 100 *µ*m^2^) at 7.8125 Hz. Optic nerve and scan field locations were recorded to reconstruct retinal positions. For higher resolution, 512 ×512 pixel images were acquired to support semi-automatic ROI detection.

### Light stimulation

For light stimulation, we used a digital light processing (DLP) projector equipped with a lightguide port (lightcrafter DPM-E4500UVBGMKII, EKB Technologies Ltd) and external UV and green light-emitting diodes (LEDs, green: 578 BP 10, F37-576; UV: 387 BP 11, F39-387; both AHF/Chroma). Visual stimuli are projected through the objective onto the retina, focusing the stimulus at the level of the photoreceptor layer (59). Both LEDs were synchronised with the microscope’s scan retrace and a bandpass filter was used to further optimise the spectral separation of mouse M- and S-opsins (390/576 dual band-pass, F59-003, AHF/Chroma). Stimulator intensity (as photoisomerisa-tion rate, 10^3^ P^∗^s^−1^ per cone) was calibrated to range from ≈0.5 (black image) to ≈ 20 for M- and S-opsins, respectively. Additionally, a steady illumination of ≈ 10^4^ P^∗^s^−1^ per cone was present during the recordings due to the 2P excitation of photopigments (56, 58).

In total, three different light stimuli were presented: (a) a full-field chirp stimulus (700 *µ*m in diameter; 23), (b) a bright moving bar (0.3× 1 mm) at 1 mm s^−1^ in eight directions, and (c) a random binary noise (“shifted” dense noise, SN) with a checker board grid of 20 × 15 checks and a checker size of 40 *µ*m at 5 Hz for 5 min, which is randomly shifted on a 10 *µ*m grid to map RFs (adapted from 33). All stimuli were centred on the recording field. Before each stimulus, the baseline was recorded for at least 20 s after the laser scanning started to avoid immediate laser-induced effects on the retinal activity (56, 58, 60).

### Immunohistochemistry

We used age-matched wild-type and *rd10* animals for the immunohistochemical imaging. Preparation of retinae was done in the same way as for functional imaging. Wholemount retinae were fixated with 4% paraformaldehyde (PFA) in 0.1 M (phosphate buffering saline) PBS for 20 minutes at 4°C. After washing the retina with 0.1 M PBS (6 × 20 minutes at 4°C) we blocked with 10% normal goat serum (NGS) and 0.3% Triton X-100 in 0.1 M PBS over night at 4°C. Afterwards the primary antibodies (Tab. 1) in a solution with 0.3% Triton X-100 and 5% NGS in 0.1 M PBS were applied and incubated for 4-9 days at 4°C. After washing again with 0.1 M PBS (6 × 20 minutes at 4°C) the samples were incubated with secondary antibodies (Tab. 2) in solution with 0.1 M PBS overnight at 4°C. After another washing session (6 × 20 minutes at 4°C) the retinae were embedded in mounting media (Vectashield, Vector Laboratories Inc., USA) on a glass slide, the dorsal side facing upwards. The samples were covered with glass cover-slips (24 × 50 mm, R. Langenbrinck GmBH, Germany), sealed with transparent nail polish and left to dry overnight at 4°C. Images were taken using a TCS SP8 confocal microscope (Leica, Germany) with a 20x (NA 0.75) oil-immersion objective and an image size of 1048 × 1048 pixels. The laser was tuned to 488 nm and 522 nm to excite STAR488 and Alexa Fluor 568, respectively. The microscope function was controlled with the LAS X software (Leica, Germany). Images were processed using the Fiji (61) software based on ImageJ (National Institutes of Health, USA, 62) and custom made python scripts.

### Data Analysis

Data analysis was organised and performed in a custom-written schema using DataJoint for Python (http://datajoint.github.io; D. Yatsenko, Tolias lab, Baylor College of Medicine; 63). Image extraction and semiautomatic region-of-interest (ROI) detection were performed using Igor Pro 8 and custom-written Python scripts.

#### Preprocessing

After the Ca^2+^ traces were extracted from individual ROIs, the raw traces were detrended by subtracting a smoothed version of the trace to remove slow drifts in the Ca^2+^ signal (60). For smoothing, we applied a Savitzky-Golay filter (64) of 3^*rd*^ polynomial order and a window length of 60 s using the Python SciPy implementation `scipy.signal.savgol_filter` (65).

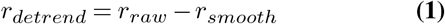

Next, the baseline activity (mean of the first 8 samples) was subtracted; the mean activity *r*(*t*) was computed, and the traces were normalised such that:

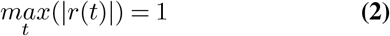

#### Quality filtering

To detect reliably responding cells, two consecutive quality filtering steps were applied. To this end, the response quality index (*QI*) was computed for moving bar (*QI*_*MB*_) and full-field chirp (*QI*_*Chirp*_) responses:

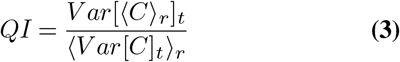

where *C* is the *T* by *R* response matrix (time samples by stimulus repetitions) and ⟨⟩ _*x*_ and *V ar*[]_*x*_ denote the mean and variance across the indicated dimension *x*, respectively. Only cells with *QI*_*MB*_ *>* 0.6 **or** *QI*_*Chirp*_ *>* 0.35 were included in the following analyses.

#### Orientation- and direction-selectivity

To detect directionselective (DS) cells, we first performed a singular values decomposition (SVD) on the normalised mean responses and projected the tuning curve on a complex exponential (*K*). The direction-selectivity index (*DSI*) was then defined as the vector length of this vector sum:

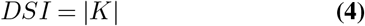

Orientation selectivity (OS) was assessed in a similar way. For details on the calculation of *DSI* and *OSI*, see (23).

#### Classification of functional RGC types

For the functional classification of RGC types, we used the method published in (26). This random forest-type classifier was trained, validated, and tested on the RGC dataset from (23). In brief, the classifier uses features of the RGC responses to the chirp and MB stimuli, the p-value of the permutation test to probe direction selectivity and soma size. The confidence score, which indicates the classification certainty, was used as a quality measure. We only included cells with a confidence score ≥ 0.25 into the RGC type-resolved analyses.

#### Receptive field estimation

We mapped RFs using the Python toolbox `RFEst` (66) using the responses to the binary shifted dense noise stimulus (see Light stimulation) We computed the temporal gradients of the Ca^2+^ signals from the detrended traces and clipped negative values:

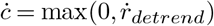

The stimulus *S*(*t*) and the clipped temporal gradients *c* were up-sampled to 10 times the stimulus frequency of 5 Hz.

Spatio-temporal RFs were computed from spline-based linear Gaussian models that were optimised with gradient descent (Adam optimiser with a learning rate of 0.1) to minimise the following loss:

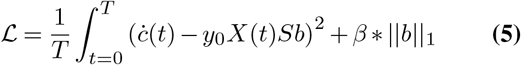

where *S* is a cubic regression spline basis, *y*_0_ is the inferred intercept, *b* are the inferred RF weights, and *β* is the weight for the L_1_-penalty on *b* to enforce sparsity in the RF.

The RF was defined as ***F*** (*x, y, τ*) = *Sb*, where *x* and *y* are the spatial dimensions and *τ* is the lag ranging from approx. 1.35 to−0.20 seconds. *S* was defined by the number of knots in space and time (*k*_*x*_, *k*_*y*_, *k*_*τ*_), corresponding to the dimensions *d* of the spatio-temporal RF (*d*_*x*_, *d*_*y*_, *d*_*τ*_) = (32, 20, 15). We optimised hyperparameters on a subset of 100 randomly drawn cells and finally set to (*k*_*x*_, *k*_*y*_, *k*_*τ*_) = (10, 12, 9) and *β* = 0.01.

Models were trained for at least 100 steps and a maximum of 2,000 steps. If the loss did not improve for 5 steps, training was stopped and the parameters resulting in the lowest loss were used as the final model.

We smoothed RFs by applying a Gaussian filter of size 5×5 pixels and a standard deviation 1 pixel to each frame. We used singular value decomposition (SVD) to decompose the RFs into a temporal *F*_*t*_(*τ*) and spatial ***F***_*s*_(*x, y*) and scaled them such that max(|*F*_*t*_|) = 1 and max(|***F***_*s*_ |) = max(|***F*** |). For each spatial RF ***F***_*s*_, we fit a 2D Gaussian ***F***_Gauss_ using the python package `astropy` (67). The area covered by two standard deviations of this Gaussian fit was used as the RF size.

Peaks in the temporal RFs were computed using `scipy.signal.find_peaks` setting the minimum peak height to 0.65 standard deviations of the temporal RFs. The main peak lag was defined as the lag of the peak with the smallest lag. To estimate the RF quality we first computed a quality metric for the SVD:

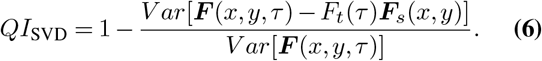

Second, we computed a quality metric for the Gaussian fit to the spatial RF:

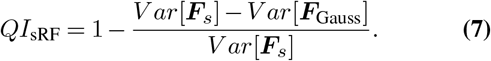

Only RFs with *QI*_SVD_ *>* 0.5, *QI*_sRF_ *>* 0.5 and main-peak-lags between 0s and 0.3s were used for the analysis.

### Statistical analysis

To quantify the differences either between cell counts (Fig. 4B-D), quality measures (Suppl. Fig. S2A,B), or RF properties (Fig. 6B, C), we performed the non-parametric Wilcoxon signed-rank test (Mann-Whitney U Test) using the Python package `scipy.stats` (v1.11.1) and employed the multicomparison Benjamini-Hochberg correction from the Python package `statsmodels.stats.multitest` (v0.14.0).

Due to large size of the dataset used for Suppl. Fig. S2, which included all imaged RGCs, we additionally calculated the effect size using the Rank-Biserial correlation (*r*_*b*_). This measure ranges from − 1 to 1, with values closer to 0 indicating a weak effect, and was calculated for the Mann-Whitney U test as follows:

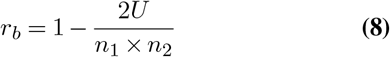

where *U* is the Mann-Whitney U statistic, *n*_1_ is the sample size of the first group, and *n*_2_ is the sample size of the second group.

To test similarities between Ca^2+^ traces, we used Pearson’s correlation (Fig. 5; Suppl. Fig. S5).

To quantify the differences in RGC type cell numbers in the *rd10* mouse line compared to wild-type, we used a binomial test (Fig. 7B). We computed the total number of cells as well as the fractions per RGC type per mouse line and age. For the binomial test, we set the test parameters *k*_*i*_ as the number of cells for each type *i* in *rd10*, and the expected proportion of cells in the corresponding RGC type, *p*_*i*_, as the proportion of cells per type *i* in wild-type. This calculation was done for P30, P45, and P90 separately. Next, we performed a two-tailed binomial test using the Python pack-age `scipy.stats` (v1.9.3). The p-value were corrected for false discovery rate (FDR, Benjamini-Hochberg); they were significant at the significance level of *α <* 0.01.

To test similarities between distributions, we calculated Jensen-Shannon-Divergence (JSD; Suppl. Fig. S6).

## Data and code availability

Data, light stimuli, as well as all custom analyses including code and notebooks to reproduce analyses will be made available at https://github.com/eulerlab upon journal publication.

## Author contributions

T.S. and T.E. designed the study with input from N.D.; N.D. performed functional imaging experiments; M.H. and N.D. performed immunohistochemistry experiments with help from T.S.; N.D. performed preprocessing; D.G., J.O., and N.D. analysed the data with the help of Y.Q.; T.S. wrote the animal protocol, N.D. and D.G. wrote the manuscript with the help from T.E., T.S., J.O., and Y.Q.

## Notes

### Competing Interest Statement

The authors have declared no competing interest.

## Bibliography

1. James K Phelan and Dean Bok. A brief review of retinitis pigmentosa and the identified retinitis pigmentosa genes. Mol Vis, 6(116):24, 2000.

2. Peter Humphries, Paul Kenna, and G Jane Farrar. On the molecular genetics of retinitis pigmentosa. Science, 256(5058):804–808, 1992.

3. Gerald J Chader. Animal models in research on retinal degenerations: past progress and future hope. Vision research, 42(4):393–399, 2002.

4. Debora B Farber, John G Flannery, and Cathy Bowes-Rickman. The rd mouse story: seventy years of research on an animal model of inherited retinal degeneration. Progress in retinal and eye research, 13(1):31–64, 1994.

5. Sascha Fauser, Janina Luberichs, and Frank Schüttauf. Genetic animal models for retinal degeneration. Survey of ophthalmology, 47(4):357–367, 2002.

6. Bo Chang, Norman L Hawes, Ron E Hurd, Muriel T Davisson, Steven Nusinowitz, and John R Heckenlively. Retinal degeneration mutants in the mouse. Vision research, 42(4): 517–525, 2002.

7. Christian P Hamel. Gene discovery and prevalence in inherited retinal dystrophies. Comptes Rendus Biologies, 337(3):160–166, 2014.

8. Rima Barhoum, Gema Martínez-Navarrete, Silvia Corrochano, Francisco Germain, Laura Fernández-Sánchez, Enrique J De la Rosa, P de La Villa, and Nicolás Cuenca. Functional and structural modifications during retinal degeneration in the rd10 mouse. Neuroscience, 155(3):698–713, 2008.

9. Theresa Puthussery, Jacqueline Gayet-Primo, Shilpi Pandey, Robert M Duvoisin, and W Rowland Taylor. Differential loss and preservation of glutamate receptor function in bipolar cells in the rd10 mouse model of retinitis pigmentosa. European Journal of Neuroscience, 29(8):1533–1542, 2009.

10. Marijana Samardzija, Hedwig Wariwoda, Cornelia Imsand, Philipp Huber, Severin R Heynen, Andrea Gubler, and Christian Grimm. Activation of survival pathways in the degenerating retina of rd10 mice. Experimental eye research, 99:17–26, 2012.

11. Claudia Gargini, Eva Terzibasi, Francesca Mazzoni, and Enrica Strettoi. Retinal organization in the retinal degeneration 10 (rd10) mutant mouse: a morphological and erg study. Journal of Comparative Neurology, 500(2):222–238, 2007.

12. Bo Chang, Norman L Hawes, Machelle Helle T Pardue, A. M. German, Ron E Hurd, Muriel T Davisson, Steven Nusinowitz, Kalpana Rengarajan, Amber P. Boyd, Sheree S Sidney, Michael Joseph Phillips, Rachael E Stewart, Rajashree Chaudhury, John M Nickerson, John R Heckenlively, and Jeffrey H Boatright. Two mouse retinal degenerations caused by missense mutations in the β-subunit of rod cgmp phosphodiesterase gene. Vision research, 47(5):624–633, 2007.

13. Enrica Strettoi, Vincenzo Pignatelli, Chiara Rossi, Vittorio Porciatti, and Benedetto Falsini. Remodeling of second-order neurons in the retina of rd/rd mutant mice. Vision research, 43 (8):867–877, 2003.

14. Steven F Stasheff. Emergence of sustained spontaneous hyperactivity and temporary preservation of off responses in ganglion cells of the retinal degeneration (rd1) mouse. Journal of neurophysiology, 99(3):1408–1421, 2008.

15. Robert E Marc, Bryan W Jones, Carl B Watt, and Enrica Strettoi. Neural remodeling in retinal degeneration. Progress in retinal and eye research, 22(5):607–655, 2003.

16. Mrinalini Hoon, Haruhisa Okawa, Luca Della Santina, and Rachel OL Wong. Functional architecture of the retina: development and disease. Progress in retinal and eye research, 42:44–84, 2014.

17. Francesca Mazzoni, Elena Novelli, and Enrica Strettoi. Retinal ganglion cells survive and maintain normal dendritic morphology in a mouse model of inherited photoreceptor degeneration. Journal of Neuroscience, 28(52):14282–14292, 2008.

18. Erika M Ellis, Antonio E Paniagua, Miranda L Scalabrino, Mishek Thapa, Jay Rathinavelu, Yuekan Jiao, David S Williams, Greg D Field, Gordon L Fain, and Alapakkam P Sampath. Cones and cone pathways remain functional in advanced retinal degeneration. Current Biology, 33(8):1513–1522, 2023.

19. Miranda L Scalabrino, Mishek Thapa, Lindsey A Chew, Esther Zhang, Jason Xu, Alapakkam P Sampath, Jeannie Chen, and Greg D Field. Robust cone-mediated signaling persists late into rod photoreceptor degeneration. Elife, 11:e80271, 2022.

20. Rachel A Care, David B Kastner, Irina De la Huerta, Simon Pan, Atrey Khoche, Luca Della Santina, Clare Gamlin, Chad Santo Tomas, Jenita Ngo, Allen Chen, et al. Partial cone loss triggers synapse-specific remodeling and spatial receptive field rearrangements in a mature retinal circuit. Cell reports, 27(7):2171–2183, 2019.

21. Joo Yeun Lee, Rachel A Care, David B Kastner, Luca Della Santina, and Felice A Dunn. Inhibition, but not excitation, recovers from partial cone loss with greater spatiotemporal integration, synapse density, and frequency. Cell reports, 38(5), 2022.

22. Jillian Goetz, Zachary F Jessen, Anne Jacobi, Adam Mani, Sam Cooler, Devon Greer, Sabah Kadri, Jeremy Segal, Karthik Shekhar, Joshua R Sanes, et al. Unified classification of mouse retinal ganglion cells using function, morphology, and gene expression. Cell reports, 40(2), 2022.

23. Tom Baden, Philipp Berens, Katrin Franke, Miroslav Román Rosón, Matthias Bethge, and Thomas Euler. The functional diversity of retinal ganglion cells in the mouse. Nature, 529 (7586):345–350, 2016.

24. J Alexander Bae, Shang Mu, Jinseop S Kim, Nicholas L Turner, Ignacio Tartavull, Nico Kemnitz, Chris S Jordan, Alex D Norton, William M Silversmith, Rachel Prentki, et al. Digital museum of retinal ganglion cells with dense anatomy and physiology. Cell, 173(5):1293– 1306, 2018.

25. Nicholas M Tran, Karthik Shekhar, Irene E Whitney, Anne Jacobi, Inbal Benhar, Guosong Hong, Wenjun Yan, Xian Adiconis, McKinzie E Arnold, Jung Min Lee, et al. Single-cell profiles of retinal ganglion cells differing in resilience to injury reveal neuroprotective genes. Neuron, 104(6):1039–1055, 2019.

26. Yongrong Qiu, David A Klindt, Klaudia P Szatko, Dominic Gonschorek, Larissa Hoefling, Timm Schubert, Laura Busse, Matthias Bethge, and Thomas Euler. Efficient coding of natural scenes improves neural system identification. PLoS computational biology, 19(4):e1011037, 2023.

27. Dominic Gonschorek, Matías A. Goldin, Jonathan Oesterle, Tom Schwerd-Kleine, Ryan Arlinghaus, Zhijian Zhao, Timm Schubert, Olivier Marre, and Thomas Euler. Nitric oxide modulates contrast suppression in a subset of mouse retinal ganglion cells. eLife, 2024. doi: 10.7554/elife.98742.1.

28. Mohamed H Farah and Stephen S Easter Jr. Cell birth and death in the mouse retinal ganglion cell layer. Journal of Comparative Neurology, 489(1):120–134, 2005.

29. Silke Haverkamp and Heinz Wässle. Immunocytochemical analysis of the mouse retina. Journal of Comparative Neurology, 424(1):1–23, 2000.

30. Brenna Krieger, Mu Qiao, David L Rousso, Joshua R Sanes, and Markus Meister. Four alpha ganglion cell types in mouse retina: Function, structure, and molecular signatures. PloS one, 12(7):e0180091, 2017.

31. Adam Bleckert, Edward D Parker, YunHee Kang, Raika Pancaroglu, Florentina Soto, Renate Lewis, Ann Marie Craig, and Rachel OL Wong. Spatial relationships between gabaergic and glutamatergic synapses on the dendrites of distinct types of mouse retinal ganglion cells across development. PloS one, 8(7):e69612, 2013.

32. Rachel A Care, Ivan A Anastassov, David B Kastner, Yien-Ming Kuo, Luca Della Santina, and Felice A Dunn. Mature retina compensates functionally for partial loss of rod photoreceptors. Cell Reports, 31(10), 2020.

33. Daniela Pamplona, Gerrit Hilgen, Matthias H Hennig, Bruno Cessac, Evelyne Sernagor, and Pierre Kornprobst. Receptive field estimation in large visual neuron assemblies using a super-resolution approach. Journal of Neurophysiology, 127(5):1334–1347, 2022.

34. Miroslav Román Rosón, Yannik Bauer, Ann H Kotkat, Philipp Berens, Thomas Euler, and Laura Busse. Mouse dlgn receives functional input from a diverse population of retinal ganglion cells with limited convergence. Neuron, 102(2):462–476, 2019.

35. Bo Chang, Norman L Hawes, Ron E Hurd, Muriel T Davisson, Steven Nusinowitz, and John R Heckenlively. A new mouse retinal degeneration (rd10) caused by a missense mutation in exon 13 of the beta-subunit of rod phosphodiesterase gene. Invest Ophthalmol Vis Sci, 41(Suppl):S533, 2000.

36. Wadood Haq, Blanca Arango-Gonzalez, Eberhart Zrenner, Thomas Euler, and Timm Schubert. Synaptic remodeling generates synchronous oscillations in the degenerated outer mouse retina. Frontiers in neural circuits, 8:108, 2014.

37. Yong Sook Goo, Kun No Ahn, Yeong Jun Song, Su Heok Ahn, Seung Kee Han, Sang Baek Ryu, and Kyung Hwan Kim. Spontaneous oscillatory rhythm in retinal activities of two retinal degeneration (rd1 and rd10) mice. The Korean journal of physiology & pharmacology: official journal of the Korean Physiological Society and the Korean Society of Pharmacology, 15(6):415, 2011.

38. Christine Haselier, Sonia Biswas, Sarah Rösch, Gabriele Thumann, Frank Müller, and Peter Walter. Correlations between specific patterns of spontaneous activity and stimulation efficiency in degenerated retina. PLoS One, 12(12):e0190048, 2017.

39. Henrike Stutzki, Florian Helmhold, Max Eickenscheidt, and Günther Zeck. Subretinal electrical stimulation reveals intact network activity in the blind mouse retina. Journal of neurophysiology, 116(4):1684–1693, 2016.

40. Ava K Bittner, Marie Diener-West, and Gislin Dagnelie. A survey of photopsias in self-reported retinitis pigmentosa: location of photopsias is related to disease severity. Retina, 29(10):1513–1521, 2009.

41. Steven F Stasheff. Clinical impact of spontaneous hyperactivity in degenerating retinas: significance for diagnosis, symptoms, and treatment. Frontiers in Cellular Neuroscience, 12:298, 2018.

42. Steven J Pittler and Wolfgang Baehr. Identification of a nonsense mutation in the rod photoreceptor cgmp phosphodiesterase beta-subunit gene of the rd mouse. Proceedings of the National Academy of Sciences, 88(19):8322–8326, 1991.

43. Dyonne T Hartong, Eliot L Berson, and Thaddeus P Dryja. Retinitis pigmentosa. The Lancet, 368(9549):1795–1809, 2006.

44. Christian Hamel. Retinitis pigmentosa. Orphanet journal of rare diseases, 1:1–12, 2006.

45. Tian Wang, Johan Pahlberg, Jon Cafaro, Rikard Frederiksen, AJ Cooper, Alapakkam P Sampath, Greg D Field, and Jeannie Chen. Activation of rod input in a model of retinal degeneration reverses retinal remodeling and induces formation of functional synapses and recovery of visual signaling in the adult retina. Journal of Neuroscience, 39(34):6798–6810, 2019.

46. You-Wei Peng, Ying Hao, Robert M Petters, and Fulton Wong. Ectopic synaptogenesis in the mammalian retina caused by rod photoreceptor-specific mutations. Nature neuroscience, 3(11):1121–1127, 2000.

47. Enrica Strettoi, Alan J Mears, and Anand Swaroop. Recruitment of the rod pathway by cones in the absence of rods. Journal of Neuroscience, 24(34):7576–7582, 2004.

48. Wan-Qing Yu, Norberto M Grzywacz, Eun-Jin Lee, and Greg D Field. Cell type-specific changes in retinal ganglion cell function induced by rod death and cone reorganization in rats. Journal of neurophysiology, 118(1):434–454, 2017.

49. Yvonne Ou, Rebecca E Jo, Erik M Ullian, Rachel OL Wong, and Luca Della Santina. Selective vulnerability of specific retinal ganglion cell types and synapses after transient ocular hypertension. Journal of Neuroscience, 36(35):9240–9252, 2016.

50. Xin Duan, Mu Qiao, Fengfeng Bei, In-Jung Kim, Zhigang He, and Joshua R Sanes. Subtypespecific regeneration of retinal ganglion cells following axotomy: effects of osteopontin and mtor signaling. Neuron, 85(6):1244–1256, 2015.

51. Maya Carleton and Nicholas W Oesch. Asymmetric activation of on and off pathways in the degenerated retina. eneuro, 11(5), 2024.

52. Robert E Marc, Bryan W Jones, James R Anderson, Krista Kinard, David W Marshak, John H Wilson, Theodore Wensel, and Robert J Lucas. Neural reprogramming in retinal degeneration. Investigative ophthalmology & visual science, 48(7):3364–3371, 2007.

53. Enrica Strettoi and Vincenzo Pignatelli. Modifications of retinal neurons in a mouse model of retinitis pigmentosa. Proceedings of the National Academy of Sciences, 97(20):11020– 11025, 2000.

54. Enrica Strettoi, Vittorio Porciatti, Benedetto Falsini, Vincenzo Pignatelli, and Chiara Rossi. Morphological and functional abnormalities in the inner retina of the rd/rd mouse. Journal of Neuroscience, 22(13):5492–5504, 2002.

55. Glenn H Kageyama and Margaret T Wong-Riley. The histochemical localization of cytochrome oxidase in the retina and lateral geniculate nucleus of the ferret, cat, and monkey, with particular reference to retinal mosaics and on/off-center visual channels. Journal of Neuroscience, 4(10):2445–2459, 1984.

56. Thomas Euler, Susanne E Hausselt, David J Margolis, Tobias Breuninger, Xavier Castell, Peter B Detwiler, and Winfried Denk. Eyecup scope—optical recordings of light stimulusevoked fluorescence signals in the retina. Pflügers Archiv-European Journal of Physiology, 457:1393–1414, 2009.

57. Kevin L Briggman and Thomas Euler. Bulk electroporation and population calcium imaging in the adult mammalian retina. Journal of neurophysiology, 105(5):2601–2609, 2011.

58. Thomas Euler, Katrin Franke, and Tom Baden. Studying a light sensor with light: multiphoton imaging in the retina. Multiphoton Microscopy, pages 225–250, 2019.

59. Katrin Franke, André Maia Chagas, Zhijian Zhao, Maxime JY Zimmermann, Philipp Bartel, Yongrong Qiu, Klaudia P Szatko, Tom Baden, and Thomas Euler. An arbitrary-spectrum spatial visual stimulator for vision research. elife, 8:e48779, 2019.

60. Klaudia P Szatko, Maria M Korympidou, Yanli Ran, Philipp Berens, Deniz Dalkara, Timm Schubert, Thomas Euler, and Katrin Franke. Neural circuits in the mouse retina support color vision in the upper visual field. Nature communications, 11(1):3481, 2020.

61. Johannes Schindelin, Ignacio Arganda-Carreras, Erwin Frise, Verena Kaynig, Mark Longair, Tobias Pietzsch, Stephan Preibisch, Curtis Rueden, Stephan Saalfeld, Benjamin Schmid, et al. Fiji: an open-source platform for biological-image analysis. Nature methods, 9(7):676–682, 2012.

62. Caroline A Schneider, Wayne S Rasband, and Kevin W Eliceiri. Nih image to imagej: 25 years of image analysis. Nature methods, 9(7):671–675, 2012.

63. Dimitri Yatsenko, Jacob Reimer, Alexander S Ecker, Edgar Y Walker, Fabian Sinz, Philipp Berens, Andreas Hoenselaar, R James Cotton, Athanassios S Siapas, and Andreas S Tolias. Datajoint: managing big scientific data using matlab or python. BioRxiv, page 031658, 2015.

64. William H Press and Saul A Teukolsky. Savitzky-golay smoothing filters. Computers in Physics, 4(6):669–672, 1990.

65. Pauli Virtanen, Ralf Gommers, Travis E. Oliphant, Matt Haberland, Tyler Reddy, David Cournapeau, Evgeni Burovski, Pearu Peterson, Warren Weckesser, Jonathan Bright, Stéfan J. van der Walt, Matthew Brett, Joshua Wilson, K. Jarrod Millman, Nikolay Mayorov, Andrew R. J. Nelson, Eric Jones, Robert Kern, Eric Larson,J J Carey, İlhan Polat, Yu Feng, Eric W. Moore, Jake VanderPlas, Denis Laxalde, Josef Perktold, Robert Cimrman, Ian Henriksen, E. A. Quintero, Charles R. Harris, Anne M. Archibald, Antônio H. Ribeiro, Fabian Pedregosa, Paul van Mulbregt, and SciPy 1.0 Contributors. SciPy 1.0: Fundamental Algorithms for Scientific Computing in Python. Nature Methods, 17:261–272, 2020. doi: 10.1038/s41592-019-0686-2.

66. Ziwei Huang, Yanli Ran, Jonathan Oesterle, Thomas Euler, and Philipp Berens. Estimating smooth and sparse neural receptive fields with a flexible spline basis. arXiv preprint 2108.07537, 2021.

67. Adrian M Price-Whelan, B. Sipőcz, H. Günther, PL Lim, SM Crawford, S Conseil, DL Shupe, MW Craig, N Dencheva, A Ginsburg, et al. The astropy project: Building an open-science project and status of the v2. 0 core package. The Astronomical Journal, 156 (3):123, 2018.

