## Supplemental Material for "Photoreceptor degeneration has heterogeneous effects on functional retinal ganglion cell types"

### Supplementary Material

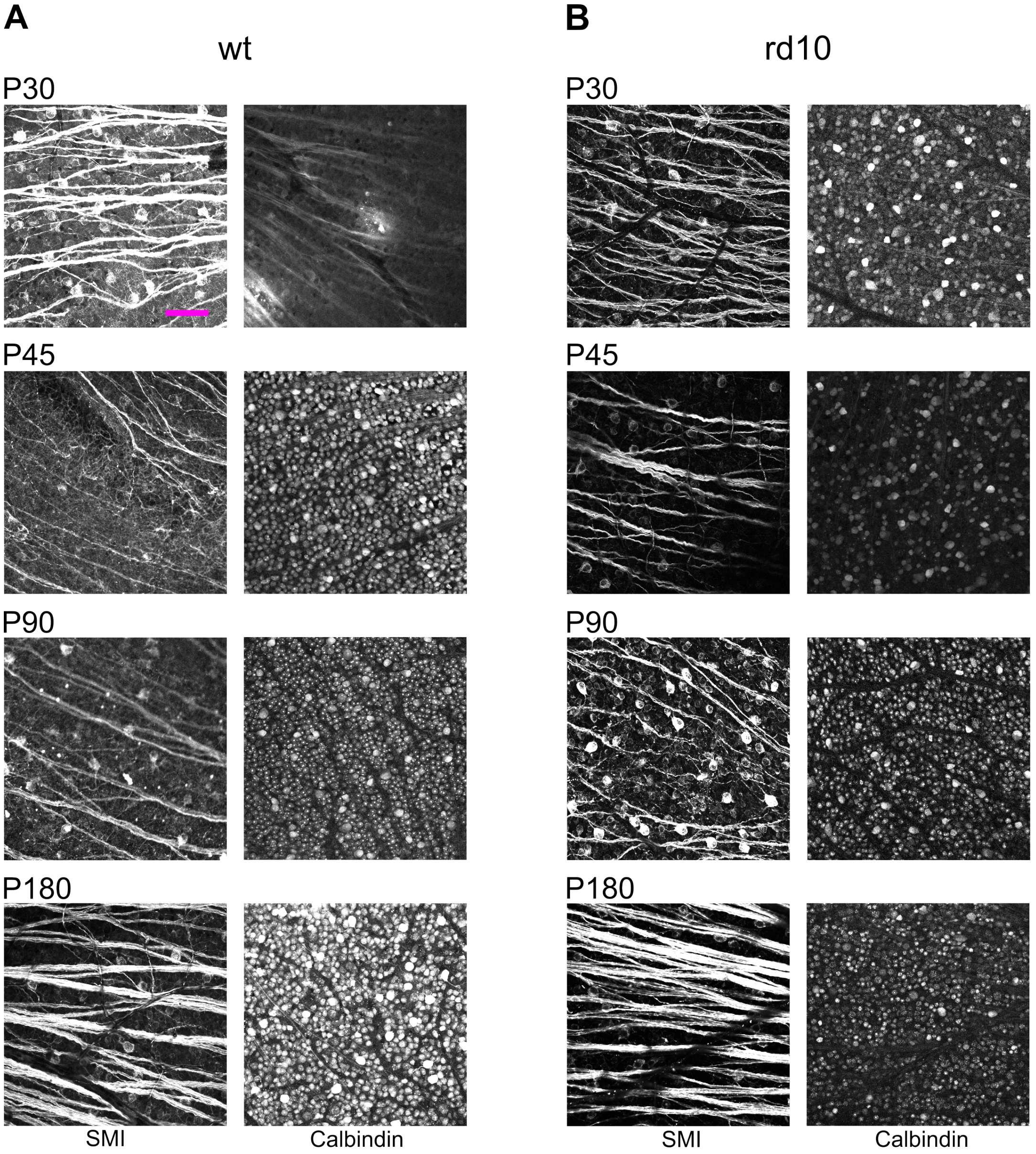


**Fig. S1. Example fields of immunohistochemistry experiments.** (**A**) Representative immunohistochemistry stainings of wild-type RGCs with SMI (left) and Calbindin (right). Rows show examples for the four selected postnatal days. Scale bar: 200$\mu$m. (**B**) Like in (A) but for *rd10* retinae.

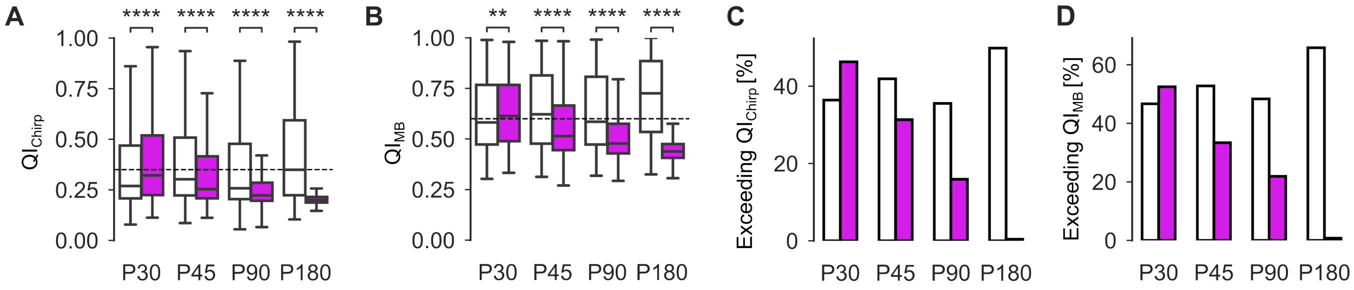


**Fig. S2. Investigating the quality of the light-evoked responses.** (**A**) Distribution of Chirp quality. Statistics for wild-type RGCs: P30 vs. P45, P90, P180: $5.28 \cdot 10^{-17}$, $1.88 \cdot 10^{-1}$, $3.66 \cdot 10^{-33}$; P45 vs. P90, P180: $3.56 \cdot 10^{-22}$, $3.58 \cdot 10^{-8}$; P90 vs. P180: $1.45 \cdot 10^{-37}$. Effect sizes for wild-type RGCs: P30 vs. P45, P90, P180: $0.115$, $-0.024$, $0.141$; P45 vs. P90, P180: $-0.138$, $0.036$; P90 vs. P180: $0.160$. Statistics for *rd10* RGCs: P30 vs. P45, P90: $1.64 \cdot 10^{-63}$, $7.85 \cdot 10^{-295}$; P45 vs. P90: $4.00 \cdot 10^{-124}$; P90 vs. P180: $2.07 \cdot 10^{-169}$. Statistics for P30, P45 vs. P180 could not be calculated due to the large differences between response quality, leading to a p-value approaching zero. Effect sizes for *rd10* RGCs: P30 vs. P45, P90, P180: $-0.178$, $-0.379$, $-0.689$; P45 vs. P90, P180: $-0.209$, $-0.551$; P90 vs. P180:$-0.360$. (**B**) Same as in (A) but for MB quality. Statistics for wild-type RGCs: P30 vs. P45, P90, P180: $1.22 \cdot 10^{-7}$, $2.08 \cdot 10^{-02}$, $1.64 \cdot 10^{-83}$; P45 vs. P90, P180: $7.36 \cdot 10^{-03}$, $5.45 \cdot 10^{-48}$; P90 vs. P180: $2.7 \cdot 10^{-63}$. Effect sizes for wild-type RGCs: P30 vs. P45, P90, P180: $0.079$, $0.023$, $0.249$; P45 vs. P90, P180: $-0.052$, $0.169$; P90 vs. P180: $0.218$. Statistics for *rd10* RGCs: P30 vs. P45, P90: $1.12 \cdot 10^{-112}$, $5.72 \cdot 10^{-283}$; P45 vs. P90: $1.52 \cdot 10^{-64}$; P90 vs. P180: $1.21 \cdot 10^{-155}$. Statistics for P30, P45 vs. P180 could not be calculated due to the large differences between response quality, leading to a p-value approaching zero. Effect sizes for *rd10* RGCs: P30 vs. P45, P90, P180: $-0.238$, $-0.376$, $-0.683$; P45 vs. P90, P180: $-0.147$, $-0.482$; P90 vs. P180:$-0.340$. (**C**) Percentage of RGCs exceeding the quality threshold for the chirp stimulus. *rd10*: P30, 46.3%; P45, 31.3%; P90, 15.9%; P180, 0.4%; wild-type: P30, 36.4%; P45, 42.9%; P90, 35.6%; P180, 49.9%. (**D**) Same as in (C) but for MB stimulus. *rd10*: P30, 52.5%, P45, 33.4%; P90, 21.9%; P180, 0.8%; wild-type: P30, 46.7%; P45, 52.8%; P90, 48.3%; P180, 65.8%; Panels (A-D) White: wild-type, magenta: *rd10*.

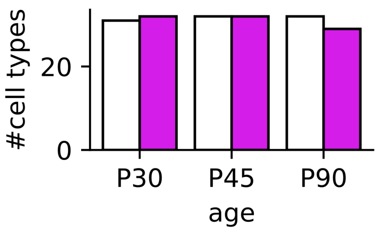


**Fig. S3. Functional cell types found in *rd10* and wild-type retinae.** Number of cell types for wild-type (white) and *rd10* (magenta) retinae at each age. Numbers for wild-type: P30/P45/P90, 31/32/32. Numbers for *rd10*: P30/P45/P90, 32/32/29.

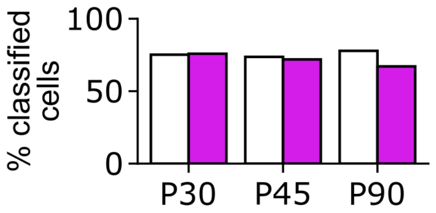


**Fig. S4. Percent of classified cells per age and mouse line.** Percent of classified cells that pass our confidence score thresholds (confidence score $\geq 0$ vs. confidence score $\geq 0.25$) (cf. Methods) for wild-type (white) and *rd10* (magenta) RGCs.

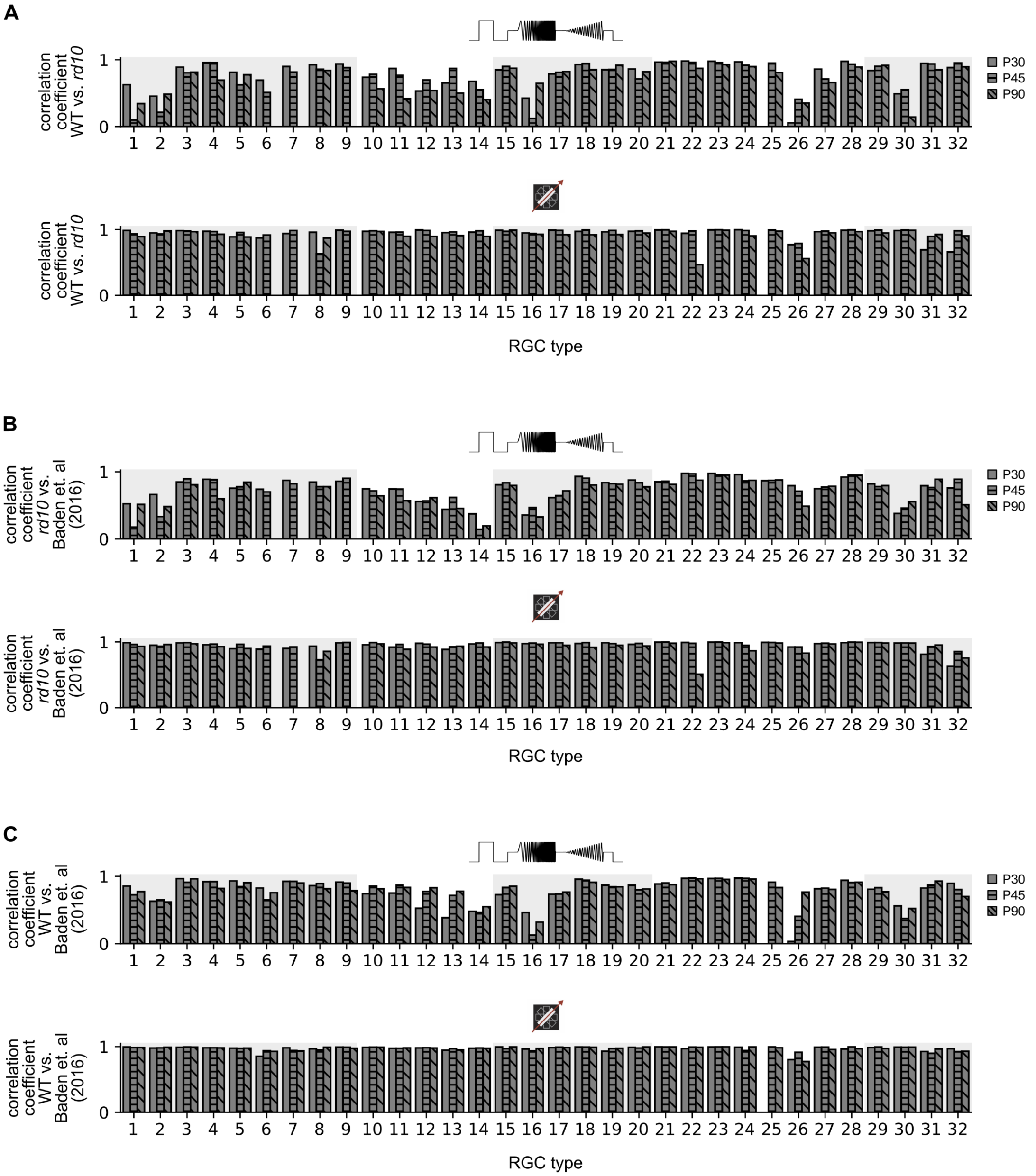


**Fig. S5. Response trace correlation for Chirp and MB stimuli.** (**A**) Correlation coefficient ($r$) between response traces of wild-type and *rd10* RGCs to chirp (top) and MB (bottom) stimuli for each cell type. P30: blank, P45: horizontal lines, P90: diagonal lines. (**B**) Same as in (A) but between response traces of *rd10* and a published RGC dataset from Baden et al. (2016). (**C**) Same as in (B) but for wild-type.

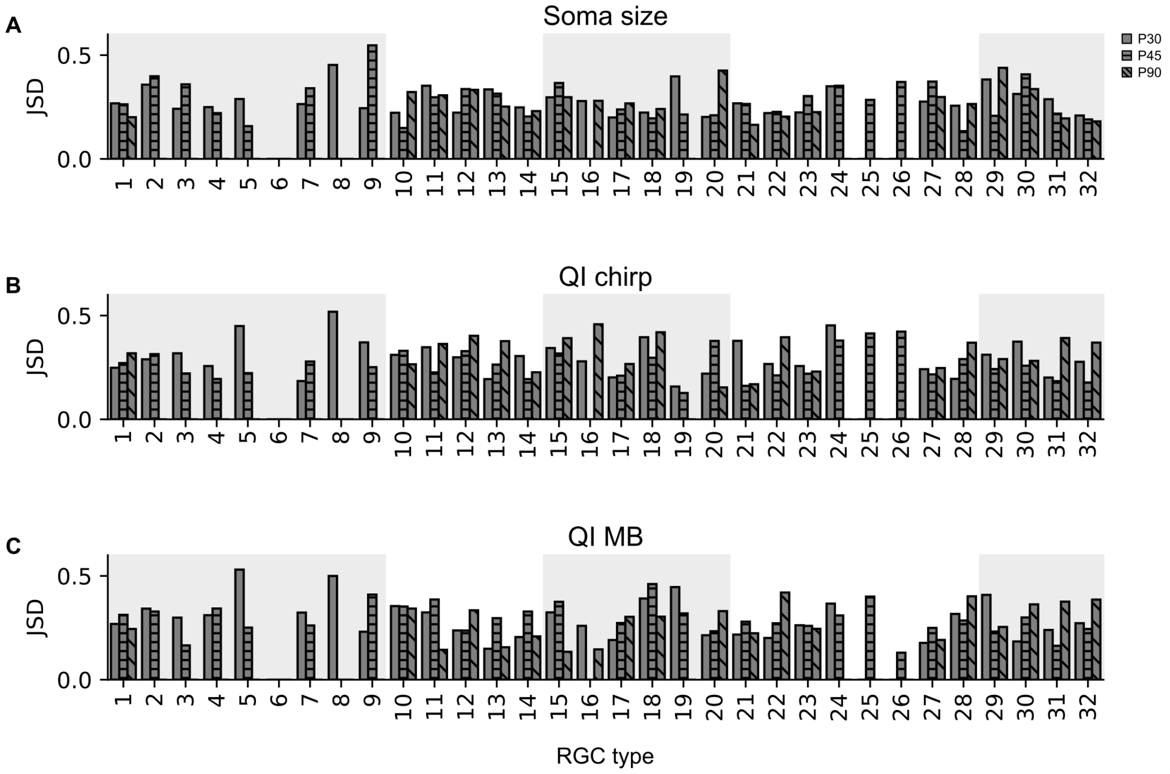


**Fig. S6. Similarity between wild-type and *rd10* distributions.** (**A**) Jensen-Shannon-Divergence (JSD, y-axis) between wild-type and *rd10* soma sizes for each cell type (x-axis). 'Empty' cell types had too few cells to compute JSD. P30: blank; P45: horizontal lines; P90: diagonal lines. (**B**) Same as in (A), but for quality distributions of chirp responses. (**C**) Same as in (A), but for quality distributions of MB responses.

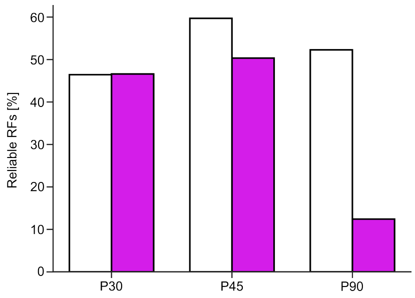


**Fig. S7. Percent of high-quality RFs per age.** Percent of RFs that passed our quality thresholds (cf. Methods) for wild-type (white) and *rd10* (magenta) RGCs.
